# A paternal cardiac lesion induces cardiac adaptation in offspring

**DOI:** 10.1101/2024.02.09.579452

**Authors:** Benedetta Coppe, María Galardi-Castilla, Andrés Sanz-Morejón, Prateek Arora, Javier Lucas, Laura Lalaguna, Enrique Lara Pezzi, Ignacio Flores, Nadia Mercader

## Abstract

**Background:** Following cardiac injury, whether the heart is permanently damaged or regenerating, distal organs are subjected to changes in physiological function. It remains largely unknown whether a cardiac lesion can affect gametes and transmit heritable changes to subsequent generations. Here, we report the influence of paternal cardiac injury on the following generation.

**Methods:** We studied the intergenerational influence of neonatal cardiac injury in the mouse, an animal model capable of regenerating the heart after early life injury. Neonatal male mice were subjected to ventricular cryoinjury, crossed at adulthood, and their sires were compared with litters derived from uninjured male mice. We used echocardiography, histology, and single nuclei RNA-sequencing to thoroughly characterize cardiac morphology, composition, function, and response to cardiac insult.

**Results:** We show that paternal cardiac injury affects the heart morphology of offspring under physiological conditions. Furthermore, in response to the same injury, the F1 generation derived from injured fathers shows better systemic and cardiac recovery, with non-pathological left ventricular enlargement and improved cardiac function during the regenerative process. This is accompanied by the activation of the immune system healing program at 3 weeks post-injury, together with enhanced transcription of genes associated with physiological hypertrophy.

**Conclusions:** The memory of a paternal neonatal lesion can be transmitted to offspring and improve their recovery from a cardiac insult.

## Introduction

Early life experiences and habits influence cardiovascular health. Physical adversities experienced early in life can increase the risk of developing cardiovascular disease (CVD) later in life (Kivimäki & Steptoe, 2018). In this regard, a study performed in zebrafish and confirmed in mice suggested that stress disrupts physiological heart development (Apaydin et al., 2020). In humans, neonatal cardiac surgery may lead to a higher incidence of long-term hypertension (Greenberg et al., 2021). Conversely, a healthy lifestyle in childhood that prevents overweight, and obesity is associated with a reduced risk of developing CVD and type 2 diabetes in adulthood (Fernandez-Jimenez et al., 2018; Laitinen et al., 2015).

*In utero* exposure to a suboptimal environment has also been associated with changes in the cardiovascular system. Maternal over-nutrition in mammals increases the risk of metabolic disorders and CVD in the offspring. For example, Blackmore et al. (2014) found that the offspring of obese mothers exhibit pathological cardiac hypertrophy associated with severe systolic and diastolic dysfunction. Likewise, exposure to a protein-restricted diet during pre- and postnatal periods in mice leads to increased reactive oxygen species production and decreased antioxidant activity in the hearts of adult offspring (Nascimento et al., 2014). The development of the cardiovascular system in newborns is influenced not only by the gestational environment, but also by the postnatal maternal diet. Indeed, maternal milk provides nutrients (*e.g.,* γ-linolenic acid) required for the cardiac metabolic switch that is essential for cardiomyocyte (CM) maturation in neonatal mice (Paredes et al., 2023).

Although prenatal maternal exposures and postnatal events are recognized to affect the cardiovascular system of the newborn later in life, it is less clear whether early parental stress experiences can have a similar impact on the cardiovascular system of the next generation. Paternal inheritance is a useful model for studying the direct germline transmission of parental experiences because, unlike maternal inheritance (which can be potentially influenced by the *in utero* environment), the male contribution to life is limited to the genetic and epigenetic material contained in the fertilizing sperm (Bohacek & Mansuy, 2017). Paternal intergenerational inheritance refers to the influence of paternal experiences on the phenotype of the F1 generation and has been observed in response to changes in diet, exposure to toxicants, or psychological trauma (Fitz-James & Cavalli, 2022). Some of these factors, together with the paternal age at conception, smoking and alcohol consumption, have also been associated with a higher risk of cardiac malformations in the offspring (Peng et al., 2019).

Interestingly, a family history of myocardial infarction (MI) influences the risk of cardiovascular disease in humans, with the risk being highest when the parental MI occurs at a younger age (Sesso et al., 2001). Whether a parental cardiac injury directly influences the offspring is not known. Results from experimentally induced chronic liver injury in rats suggest that organ damage may indeed lead to positive adaptation to the same insult in the offspring (Zeybel et al., 2012). We performed acute cardiac injury by cryo-lesion of the left ventricle in neonates, and found molecular, anatomical, and functional cardiac effects in their offspring, providing evidence for intergenerational consequences of a paternal history of cardiac injury in mammals.

## Results

### Paternal cardiac injury induces temporal systemic changes in offspring in response to cardiac injury

We used ventricular cryoinjury (CI) to study the intergenerational effect of early-life cardiac injury and healing. Parental generation male mice underwent CI at postnatal day 1 (P1), were raised to adulthood, and mated with control females. All animals were fertile and produced offspring. As a control, uninjured male mice from the same genetic background and age were used and similarly crossed with females from the same stock as the experimental group. Animals of the F1 generation were left uninjured or subjected to CI at P1 (Figure 1A). Hereinafter, we use acronyms to refer to the F1 generation: “P” represents “paternal generation”, “F” the “F1 generation”, “u” stands for “uninjured”, and “i” for “injured”. Thus, the uninjured offspring of uninjured fathers are termed “Pu-Fu”, the uninjured offspring of injured fathers are termed “Pi-Fu”, and the injured offspring of uninjured or injured fathers are termed “Pu-Fi” or “Pi-Fi”, respectively (Figure 1A).

**Figure 1.**
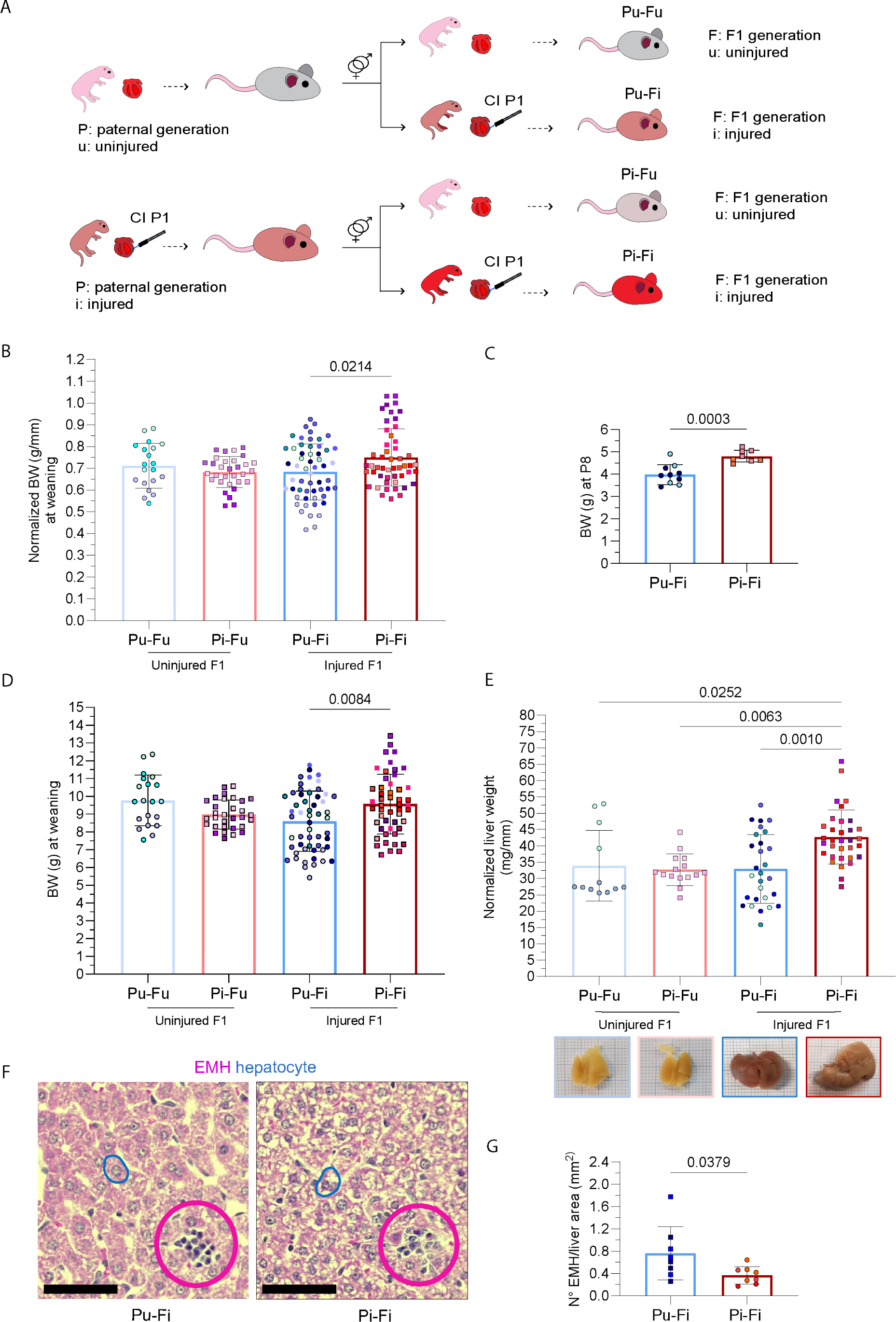
Systemic changes in the offspring of uninjured or injured fathers at weaning. **A,** Scheme of the experimental design utilized to investigate the effects of paternal heart injury on the next generation. In the control group, fathers were left uninjured; in the experimental group, fathers underwent cardiac cryoinjury at P1. Offspring from both groups were left uninjured or injured at P1 and raised. P, paternal generation; F, F1 generation; u, uninjured; i, injured; CI, cryoinjury. **B,** Body weight (BW) of F1 mice (g) normalized to tibial length (mm) under physiological condition and after CI at weaning (3 weeks *post-partum* (wpp). Each bar represents data from siblings of multiple sires, identified by different colors (number of sires: Pu-Fu=3, Pi-Fu=4, Pu-Fi=8, Pi-Fi=8). Shown is mean and S.D. One-way ANOVA, Tukey’s multiple comparison test. **C,** Injured offspring BW (g) at P8. Each column shows siblings from two sires. Shown is mean and S.D. Unpaired t-test. **D,** F1 BW (g) under physiological condition and after cardiac injury at weaning (3 wpp). Each column represents data from siblings of multiple sires, identified by different colors (number of sires: Pu-Fu=3, Pi-Fu=4, Pu-Fi=8, Pi-Fi=8). Shown is mean and S.D. Kruskal-Wallis test, Dunn’s multiple comparisons test. **E,** Liver weight (mg) normalized to tibial length (mm) at weaning (3 wpp). Each bar represents data from siblings of multiple sires, identified by different colors (number of sires: Pu-Fu=2, Pi-Fu=2, Pu-Fi=6, Pi-Fi=7). Shown is mean and S.D. Kruskal-Wallis test, Dunn’s multiple comparisons test. On the bottom, representative pictures of livers from uninjured (after 10% Formalin fixation) and injured F1 (right after necropsis). **F,** Hematoxylin/eosin staining of liver sections from injured F1 mice at weaning (3 wpp). Blue lines enclose hepatocytes containing or not glycogen (stained in white). The Pu-Fi section is representative of 5 out of 8 analyzed samples, Pi-Fi of 7 out of 8 samples. Fuchsia circles enclose extramedullary hematopoiesis (EMH) *loci*. Scale bar: 50 µm. **G,** Quantification of EMH sites normalized to liver area (mm^2^) in injured F1 mice at weaning (3 wpp). Each bar represents data from siblings of multiple sires, identified by different colors (number of sires: Pu-Fi=2, Pi-Fi=2). Shown is mean and S.D. Mann-Whitney test.

At weaning, 3 weeks *post-partum* (wpp), the survival of Pu-Fi and Pi-Fi was comparable (Figure S1A). Pu-Fi and Pi-Fi size, evaluated as tibial length, were similar but smaller than the uninjured pups (Figure S1B), suggesting a delayed growth in response to the injury. However, Pi-Fi mice showed an increased normalized BW (BW *vs* tibial length) compared to Pu-Fi (Figure 1B), due to the increased body weight (BW) gain observed at P8 (Figure 1C) and at weaning (Figure 1D). This phenotype was more pronounced in males (Figure S1C) than in females (Figure S1D).

Necropsy of the four groups at weaning revealed a difference in normalized liver weight between Pi-Fi and the other groups. Pi-Fi mice had heavier livers (Figure 1E) in line with the BW gained. Analysis of liver tissue in the Pu-Fi and Pi-Fi groups revealed decreased glycogen storage in 5 out of 8 Pu-Fi livers compared with 1 out of 8 Pi-Fi livers (Figure 1F). Both groups showed sites of extramedullary hematopoiesis (EMH), which occurs physiologically in mouse fetal liver (Sonoda & Sasaki, 2012). EMH is no longer detectable approximately two weeks after birth but is reactivated in response to stressors such as infection, anemia, and MI (Alamo et al., 2017). The significant difference in the number of EMH *loci* between the Pu-Fi and Pi-Fi groups (Figure 1G) indicated a reduced extramedullar hematopoietic demand or a faster recovery in the offspring of injured fathers. Overall, the increase in body and liver weights in Pi-Fi at weaning suggests a systemic protective response to cardiac injury in the offspring of injured fathers.

Analysis of the same parameters in F1 mice at adulthood revealed that body (Figure S1E) and liver (Figure S1F) weights were similar between the uninjured and injured groups, suggesting that the systemic changes observed in response to cardiac injury in Pi-Fi mice are transient and limited to early life.

### Cardiac changes can be observed in the offspring of injured fathers under physiological conditions and in response to cardiac injury

We next asked whether the paternal cardiac injury might also affect the morphology and functionality of the offspring heart. We evaluated heart weight in the F1 generation under physiological conditions, normalized by tibial length to account for the effect of the litter size on pup heart dimensions (Figure S2A). While no increase in heart weight of Pi-Fu mice with respect to Pu-Fu mice was evident at weaning (Figure S2B,C,D), echocardiographic analyses revealed a significantly heavier (*i.e.* normalized left ventricle (LV) mass corrected) and larger (*i.e.* volume) LV in Pi-Fu mice during both systole and diastole (Figure 2A,B). At the functional level, the E velocity was modestly but significantly higher in Pu-Fi mice than in Pu-Fu mice (Figure 2A,B), with no differences in other cardiac functional parameters and wall thickness between the two groups (Figure S2E).

**Figure 2.**
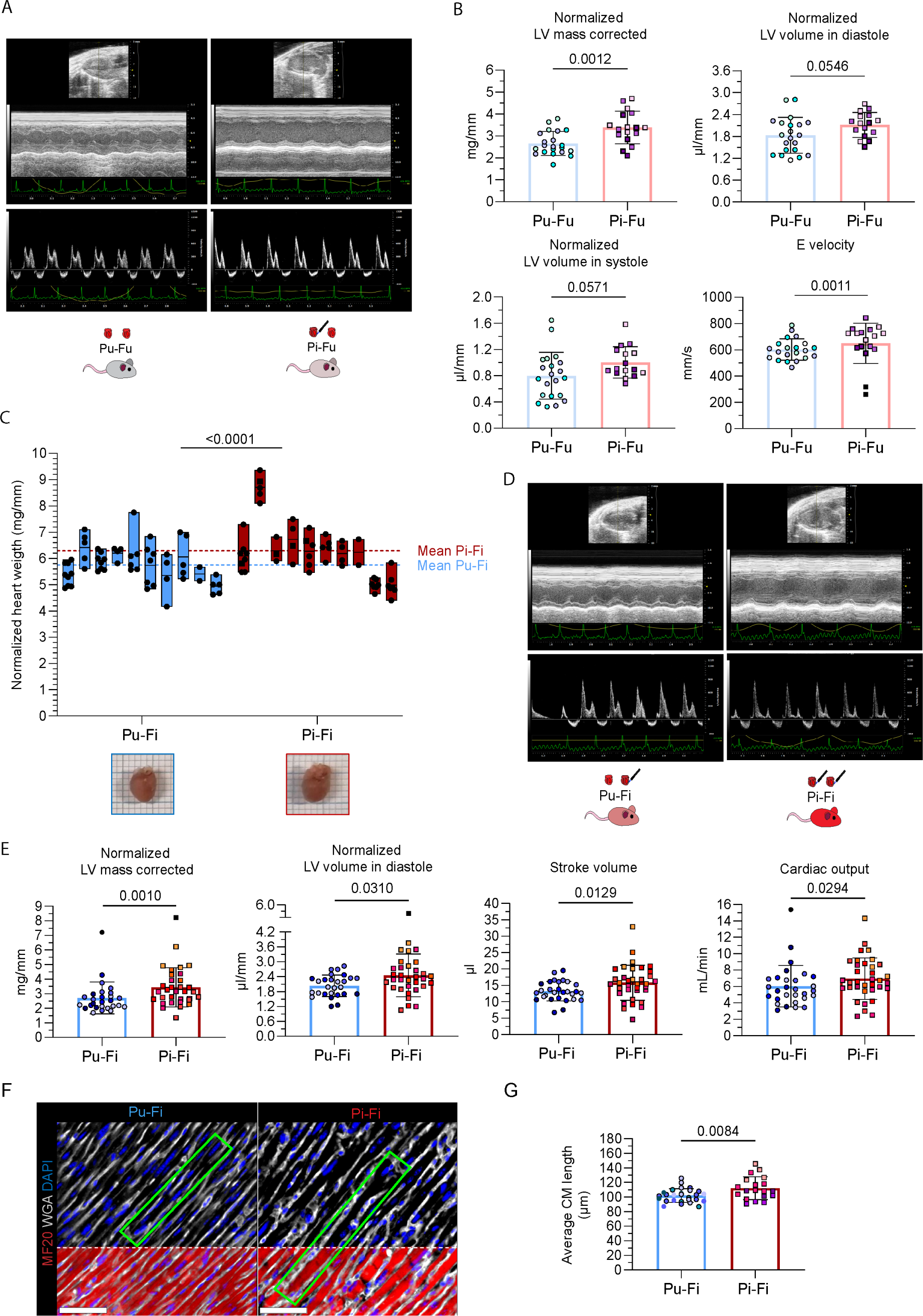
Cardiac morphological and functional changes in the offspring of uninjured or injured fathers at weaning. **A,** Representative echocardiographic images of hearts from uninjured mice of uninjured and injured fathers 3 weeks *post-partum* (wpp) (from top to bottom: two dimensional (2D), M-mode (MM), and pulse wave Doppler (PW) echocardiography). **B,** Morphological and functional assessment of cardiac features of the F1 mice of uninjured and injured fathers under physiological conditions at weaning by echocardiography. Graphs show corrected LV mass (unpaired t-test), LV volume in systole (unpaired t-test) and diastole (Mann-Whitney test), and mitral valve E velocity (unpaired t-test) at 3 wpp. Morphological data are normalized to tibial length. Dots represent individual hearts from different sires, identified by colors (n = 3 Pu-Fu, n = 3 Pi-Fu). Shown is mean and S.D. **C,** Normalized heart size of siblings from different litters represented as floating bars with line at mean. Each bar represents siblings of a single family. Dots represent individual animals. Two-way ANOVA. **D,** Representative echocardiographic images of hearts from injured mice of uninjured and injured fathers 3 wpp (from top to bottom: 2D, MM, and PW echocardiography). **E,** Morphological and functional assessment of cardiac features of the injured F1 mice of uninjured and injured fathers at weaning (3 wpp) by echocardiography. Graphs show corrected LV mass (Mann-Whitney test), LV volume in diastole (unpaired t-test), stroke volume (Mann-Whitney test), and cardiac output (unpaired t-test). Morphological data are normalized to tibial length. Dots represent individual hearts from different sires, identified by colors (number of sires: Pu-Fi=5, Pi-Fi=6). Shown is mean and S.D. **F,** Representative immunofluorescence of longitudinal plane CMs stained with MF20 (anti-myosin heavy chain, red) WGA (wheat germ agglutinin, white) to highlight membranes, and DAPI (blue) to stain nuclei. Scale bar: 50 µm. **G**, Quantification of average CM length at 3 wpi. Each bar represents data from siblings of multiple sires, identified by different colors (number of sires: Pu-Fi=4, Pi-Fi=4). Shown is mean and S.D. Unpaired t-test. P, paternal generation; F, F1 generation; u, uninjured; I, injured; LV, left ventricle; CM, cardiomyocyte.

We then compared the cardiac morphology and functionality in the injured F1 mice of uninjured and injured fathers at weaning. The normalized heart weight in injured F1 mice was significantly greater than that of Pu-Fi mice in 8 out of 10 Pi-Fi litters (Figure 2C). This phenotype was observed in both sexes but was more pronounced in males (Figure S3A) than in females (Figure S3B). Similarly, the normalized LV corrected mass was significantly greater (P=0.0010) in Pi-Fi mice than in Pu-Fi mice (Figure 2D,E). These findings suggest that cardiac remodeling after injury results in larger hearts in the offspring of injured fathers. Functionally, echocardiography revealed similar ejection fraction and fractional shortening between groups (Figure S3C); however, stroke volume and cardiac output were both significantly higher in the Pi-Fi group than in the Pu-Fi group (19.3% and 22.1%, respectively) (Figure 2D,E). The increased LV dimension in Pi-Fi mice in the absence of wall thickening (*i.e.* interventricular septum and LV posterior wall; Figure S3C), together with the increased hemodynamic measures, are signs of eccentric hypertrophy (Mihl et al., 2008; Müller & Dhalla, 2013). By immunostaining the sarcolemma, we observed that CM length was slightly but significantly higher (11.5%) in Pi-Fi mice than in Pu-Fi mice (Figures 2F,G), indicating cardiac remodeling. In sum, the offspring of injured fathers exhibit congenital changes in heart morphology and function that are exacerbated in response to injury, possibly leading to a more compliant LV and increased cardiac functionality.

We next assessed the persistence of the cardiac phenotype and observed that cardiac remodeling was still evident in adulthood when comparing injured F1 mice with uninjured F1 mice (Figure S4A), but no differences were detected between Pu-Fu and Pi-Fu (Figure S4B) nor between Pu-Fi and Pi-Fi (Figures S4C,D). Thus, the “memory” of a paternal cardiac injury transiently influences neonatal cardiac development and its response to an early-life insult.

### Early-life cardiac changes in the offspring of injured fathers are partially induced by paternal thoracotomy

To understand the mechanism involved in the transmission of paternal cardiac damage to F1 mice, we independently tested the effects of paternal cryoanesthesia and sham surgery, both steps performed during the CI surgery, in the next generation. We anesthetized the fathers at P1 by deep hypothermia (here named “cold”) or performed a thoracotomy (*i.e.,* sham), consisting of opening of the chest without damaging the heart. We then compared the response to CI in the F1 generation (hereafter named “Pc-Fi”, where “c” stands for “cold”; and Ps-Fi, where “s” stands for “sham”) (Figure 3A).

**Figure 3.**
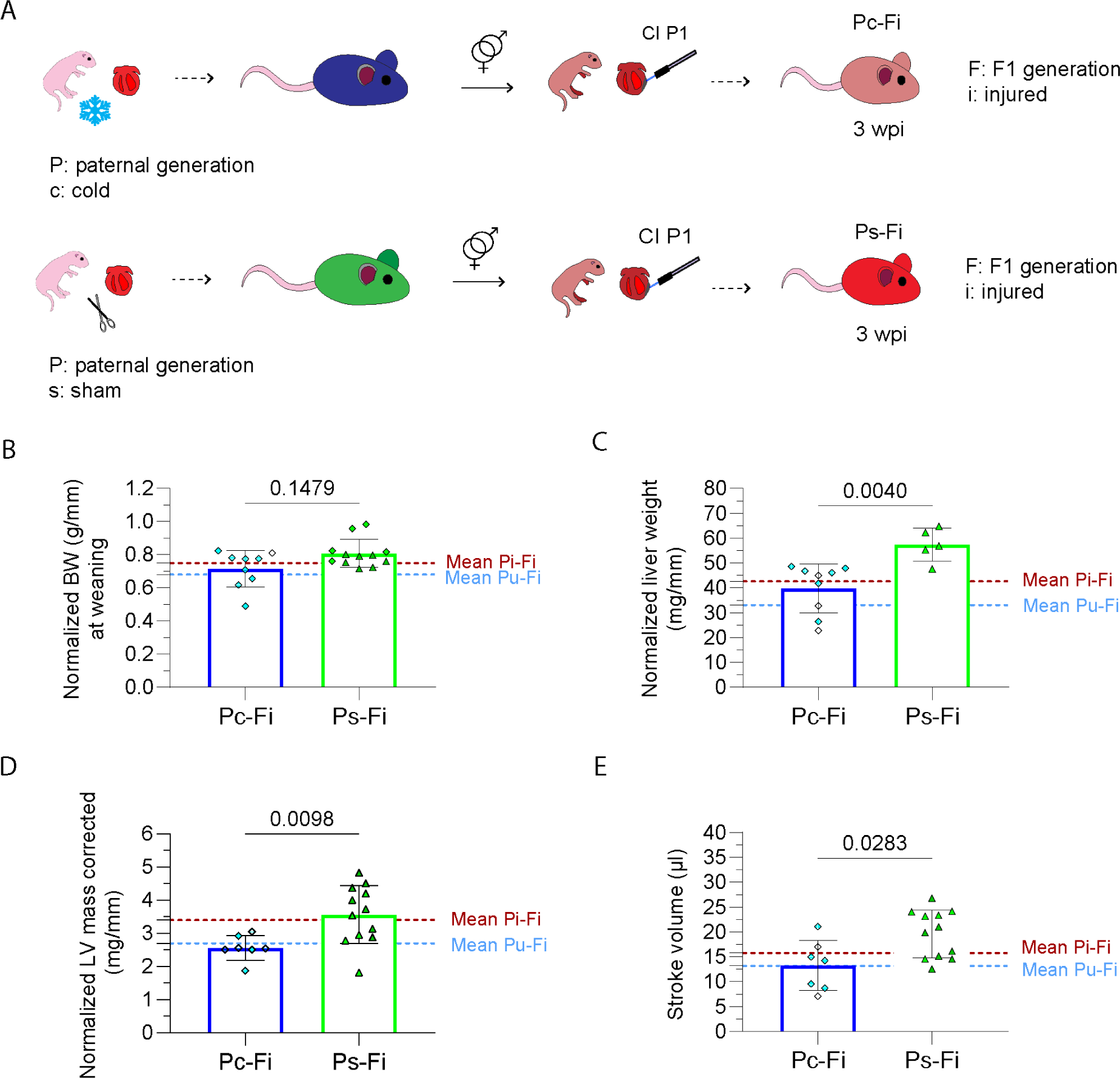
Effects of paternal hypoxia/hypothermia and sham surgery in the offspring at weaning. **A,** Schematic representation of experiments to investigate the effects of paternal deep hypothermia (“cold”) and sham surgery. Fathers underwent cryoanesthesia or thoracotomy at P1 and their offspring were injured (CI of the heart) at P1. P, paternal generation; F, F1 generation; u, uninjured; i, injured; c, cold; s, sham; CI, cryoinjury. **B,** Scatter dot plot showing body weight (BW) normalized to tibial length (g/mm) of the injured offspring of “cold” and “sham” fathers at weaning (3 weeks *post-partum* (wpp)). Shown are the values in the offspring produced by three sires (one “cold” and two “sham” fathers), identified by different colors. Dotted red and cyan lines represent the mean value in Pi-Fi and Pu-Fi mice, respectively. Mann-Whitney test. **C,** Plot showing normalized liver weight (g/mm) at weaning (3 wpp) of the injured offspring of “cold” and “sham” fathers. Shown are the values of the offspring produced by three sires (one “cold” and two “sham” fathers), identified by different colors. Dotted red and cyan lines represent the mean value in Pi-Fi and Pu-Fi mice, respectively. Mann-Whitney test. **D,** Scatter dot plot showing normalized LV mass corrected (mg/mm) of the injured offspring of “cold” and “sham” fathers at weaning (3 wpp). Shown are the values of the offspring produced by three sires (one “cold” and two “sham” fathers), identified by different colors. Dotted red and cyan lines represent the mean value in Pi-Fi and Pu-Fi mice, respectively. Mann-Whitney test. **E,** Plot showing stroke volume of the injured offspring of “cold” and sham fathers at weaning (3 wpp). Shown are the values of the offspring produced by three sires (one “cold” and two “sham” fathers), identified by different colors. Dotted red and cyan lines represent the mean value in Pi-Fi and Pu-Fi mice, respectively. Mann-Whitney test.

In response to CI, normalized BW (Figure 3B), normalized liver weight (Figure 3C), heart weight (Figure 3D), and stroke volume (Figure 3E) were all higher in Ps-Fi mice than in Pc-Fi mice at weaning. Notably, whereas the paternal neonatal hypoxic/hypothermic state induced in the next generation a similar injury response to that observed in F1 mice of uninjured fathers (*i.e.* Pc-Fi and Pu-Fi) (Figures 3B,D,E), the injured offspring of sham-operated fathers partially resembled the offspring of injured fathers (*i.e.* Ps-Fi and Pi-Fi) (Figures 3B,D). As in CI, thoracotomy can induce sterile inflammation, leading to the formation of pericardial adhesions, which in turn compromises cardiac functionality (Bailey et al., 1984; Jiamsripong et al., 2010) and alters cardiac gene expression (Botos et al., 2023). Overall, these data show that neonatal surgery, but not deep hypothermia, influences cardiac function as well as body and liver weight in the next generation.

### Offspring of injured fathers show changes in cell type composition and upregulation of the inositol triphosphate/calcium signaling pathway following neonatal cardiac injury

To better understand the causes of the cardiac changes observed in the hearts of Pi-Fi mice at weaning, we further investigated their cell composition and gene expression changes using single nuclei RNA-sequencing (snRNA-seq). At 3 wpp, we collected the whole heart from two Pu-Fi mice and two Pi-Fi mice and processed the samples for snRNA-seq. We captured 35,458 nuclei from the four samples and sequenced a median of at least 1240 genes per nucleus.

Sequencing allowed us to identify cell clusters based on known marker genes from the literature (see supplementary methods and Figures S5A, B). The cell clusters for cardiomyocytes, fibroblasts, T cells, and other immune cells were then subjected to individual subclustering (Figures 4A and S6A-D). Inspection of the proportions of all cell populations between Pu-Fi and Pi-Fi samples revealed that adipocytes, lymphocytes, and epicardial cells were more abundant in Pi-Fi samples, whereas myofibroblasts were lower in abundance (Figure 4B). Indeed, adipocytes were almost 20 times more abundant in Pi-Fi samples than in Pu-Fi samples (284 *vs* 15 nuclei) and showed the largest difference between the two groups. Among T cell populations, we identified natural killer T cells (NKT), cytotoxic T cells (T8), and regulatory T cells (Treg) to be present in greater abundance in Pi-Fi samples. Similarly, fibroblasts expressing genes involved in extracellular matrix organization (“FBs_2”), and M2a/c macrophages, likely involved in wound healing and remodeling (see supplementary methods), were higher in Pi-Fi samples. Other cell populations were observed as generally more abundant in Pi-Fi samples; however, the changes were not markedly different across the replicates, with “Pu-Fi 1” samples being similar to Pi-Fi samples.

**Figure 4.**
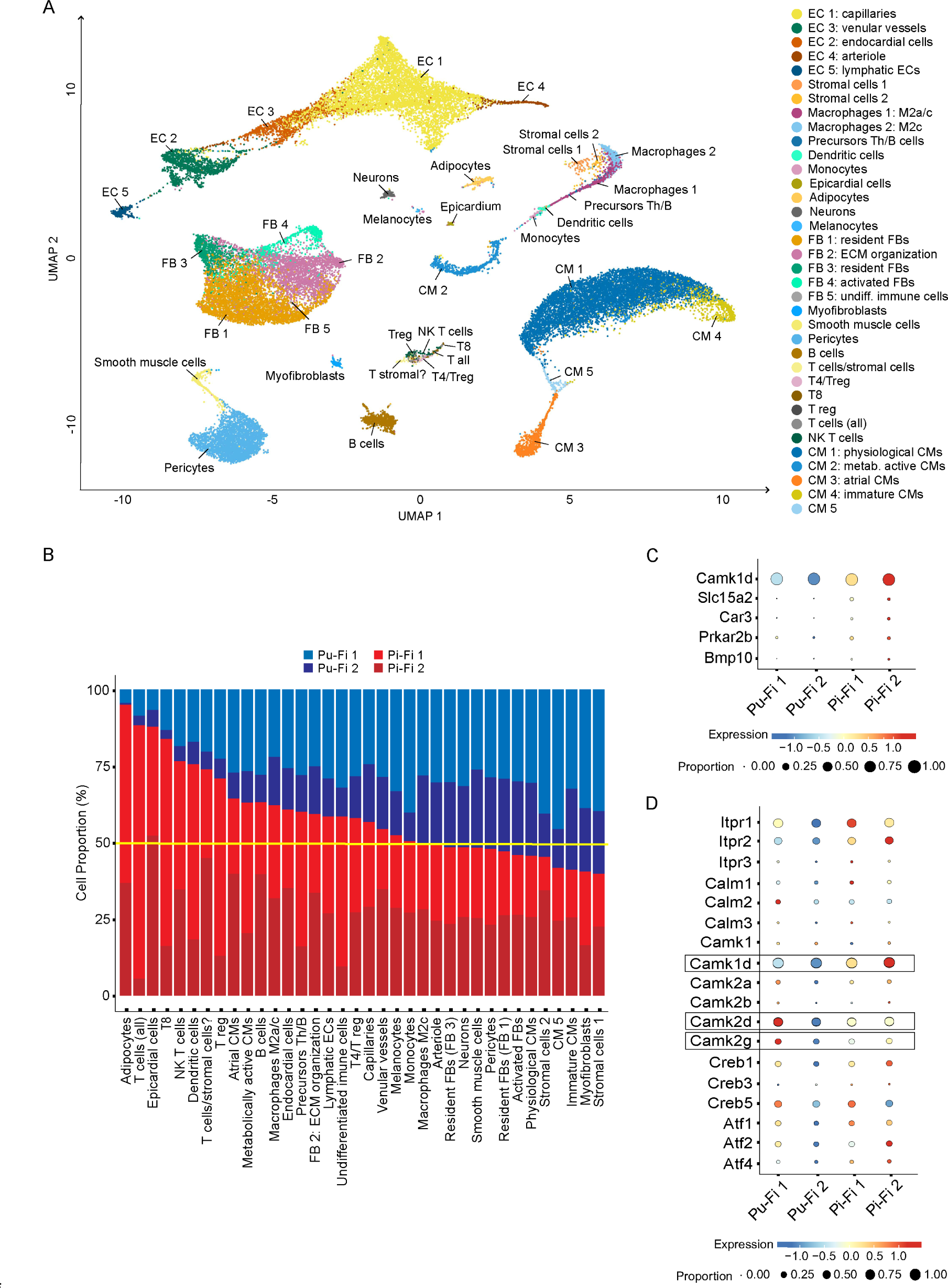
Whole heart snRNA-seq of injured offspring at weaning. **A,** UMAP visualization of unsupervised clustering of 21-dpi whole-heart samples from two replicates per condition (F1 from uninjured *vs* injured fathers). The legend indicates the identified cell populations. EC, endothelial cell; FB, fibroblasts; T reg, T regulatory cells; Th, T helper cells, NK, natural killer cells; CM, cardiomyocytes, undiff., undifferentiated; metab., metabolically. **B,** Cell proportion representation of identified cell populations. P, paternal generation; F, F1 generation; u, uninjured; i, injured. **C,** Bubbleplot of selected genes among those differentially expressed in multiple cell populations identified upon pseudobulk analysis. Dot color indicates gene expression levels scaled across samples (*i.e.* whole heart); dot size denotes proportion of cells expressing the gene. **D,** Bubbleplot of genes belonging to the inositol triphosphate/calcium signaling pathway. Dot color indicates gene expression levels scaled across samples; dot size denotes proportion of cells expressing the gene.

We then used pseudobulk-EdgeR-LRT (Squair et al., 2021) to search for differentially expressed genes (DEGs) across all cell populations. We identified 157 DEGs between Pu-Fi and Pi-Fi samples (Table S1). We focused on those genes differentially expressed across multiple cell populations, which would imply a broad and coordinated response to cardiac injury (Figure S6E). We found that *Camk1d*, a calcium/calmodulin-dependent protein kinase, and *Slc15a2* (solute carrier family 15 member 2), a transporter responsible for the absorption of small peptides, were more highly expressed in Pi-Fi samples in most of the cell populations (Figures 4C and S6E). We next asked whether any of the pathways linked to these genes were also activated. We found that the phosphoinositol-calcium (Pi3-Ca^2+^) pathway, associated with *Camk1d,* showed an overall upregulation of multiple pathway genes. The Pi3-Ca^2+^ pathway is physiologically implicated in cardiac hypertrophic growth and increased cardiac contractility (Ghigo et al., 2017). Notably, while the more commonly described member *Camk2d* (Dewenter et al., 2017) was highly expressed in Pu-Fi samples, *Camk1d* was highly differentially expressed between the two groups (Figure 4D).

We further identified the upregulated expression of several other genes in Pi-Fi samples, including *Car3*, encoding carbonic anhydrase III and previously described to be activated in hypertrophic response to adrenergic stimulation (Alvarez et al., 2007); *Prkar2b* (protein kinase CAMP-dependent type II regulatory subunit beta), a gene likely involved in the onset of cardiac hypertrophy (Saad et al., 2018); and *Bmp10* (bone morphogenetic protein 10), described to have a role in cardiac proliferation and repair after MI (Sun et al., 2014) (Figure 4C), among others (Figure S6E). To explore whether the enhanced *Bmp10* expression translated to a superior regenerative ability of the heart, we compared the regenerative response in Pu-Fi and Pi-Fi mice by measuring CM proliferation at 7 days post-injury and scar resolution at 3 wpp. No differences were detected in either parameter at these time points (Figures S7A-D).

Overall, snRNA-seq analysis of whole hearts indicated an increase in adipocytes, T lymphocytes, M2 macrophages and fibroblasts involved in extracellular matrix remodeling and scar healing in the offspring of injured fathers with respect to the offspring of uninjured fathers, and a decrease in myofibroblasts. These changes suggest the enhanced involvement of the immune system in the healing process in the offspring of injured fathers. Also, the higher levels of genes involved in the Pi3-Ca^2+^ pathway in Pi-Fi mice support the phenotypic observation of cardiac hypertrophic response to cardiac damage.

## Discussion

The intergenerational impact of early-life cardiac dysfunction remains unknown. Here, we used a neonatal mouse model of transient cardiac injury to examine its effects on the next generation. We show that the offspring of injured fathers develop a protective cardiac phenotype in response to the same injury, characterized as a non-pathological remodeling of the heart involving protective cell populations and improved cardiac functionality.

Paternal neonatal thoracotomy but not cryoanesthesia affected the offspring heart in a manner similar to paternal CI. This suggests that the sterile inflammation, pericardial adhesion and possibly cardiac dysfunction (Bailey et al., 1984; Jiamsripong et al., 2010), which occur in both CI and thoracotomy, but not following hypoxia/hypothermia, may affect male gametes and the next generation.

Under physiological conditions, the offspring of injured fathers exhibited transient enhanced cardiac growth at weaning, evidenced by increased LV dimensions and mitral valve E velocity. Considering the young age of the mice when this occurred, and its transient nature (unchanged in adults), we hypothesize that this is likely related to a physiological reduction in LV diastolic pressure and faster LV relaxation early in life. Interestingly, the increased LV dimensions under physiological conditions were more pronounced after injury, possibly due to the increased stroke volume and cardiac output, which would lead to the establishment of mild eccentric hypertrophy. The changes in gene expression in the PI3/Ca^2+^ pathway, which is involved in physiological cardiac hypertrophic growth and increased cardiac contractility after MI (Ghigo et al., 2017), were consistent with this phenotype. Among the calcium/calmodulin-dependent kinase family involved in the PI3/Ca^2+^ pathway, we found increased expression of *Camk1d* in the offspring of injured fathers, whereas the offspring of control fathers preferentially expressed the canonical *Camk2d* (Dewenter et al., 2017). In the setting of ischemia/reperfusion injury, Camk2 appears to stimulate a pro-inflammatory response and is associated with scar formation (Gray et al., 2017). The cardiac role of Camk1d is not known, but a whole-body knockout mouse model for this protein manifests reduced heart weight (Groza et al., 2023). The increase in *Camk1d* expression accords with the increase in the weight of injured hearts of the offspring of injured fathers. We hypothesize that the higher expression of *Camk1d* might be linked to a physiological hypertrophic response that maintains a functionally enlarged heart, similar to athletes and pregnant women (Toncelli et al., 2022). Other genes found to be expressed at higher levels in the injured offspring of injured fathers further support the activation of hypertrophic and protective mechanisms.

Single-nuclei transcriptomics of whole hearts at weaning allowed us to further investigate whether the memory of paternal cardiac injury triggers the involvement of different cells in response to the same injury in the F1 generation. The most striking difference was the increase in adipocyte nuclei in the offspring of injured fathers. In mice, fat depots are located sub-epicardially in the atrial-ventricular groove of the heart. They correspond to human epicardial adipose tissue (Yamaguchi et al., 2015) and resemble brown and beige adipocytes (Sacks et al., 2013). In humans, epicardial adipose tissue has been described as a source of cardiac heat and protection of the myocardium under ischemia or hypoxia (Lacobellis, 2015). Interestingly, the number of epicardial cells was also higher in the offspring of injured fathers. Because epicardial cells proliferate rapidly in response to injury (Darehzereshki et al., 2015), one possibility is that epicardial cell activation is more enhanced in injured offspring of injured *versus* uninjured fathers and is accompanied by their differentiation into adipocytes (Yamaguchi et al., 2015).

The injured offspring of injured fathers also displayed an enhanced immune response, with more abundant NKT, T8, and T-reg nuclei. NKT cells are involved in cardiac remodeling following MI and have an anti-inflammatory and protective function (Sobirin et al., 2012) by activating other immune cells, such as dendritic cells, monocytes, macrophages, T and B cells (van Puijvelde & Kuiper, 2017), which may explain the increase observed in Pi-Fi mice. T-reg cells have multiple roles in cardiac tissue repair: they promote macrophage polarization towards the anti-inflammatory and healing M2 type, reduce the expression of α-SMA in fibroblasts, and improve cardiac function in ischemic heart tissue (Kino et al., 2020). The evident differences in the abundance of T-reg cells together with the increased number of macrophages involved in wound healing and remodeling, the reduced presence of myofibroblasts and the increased cardiac output, suggest an overall better healing in the offspring of injured fathers. Further studies evaluating injuries performed at P7, a time window in which natural regeneration is lost (Porrello et al., 2011), might provide opportunities to explore whether a paternal lesion not only allows better recovery after injury, but also prolongs the regeneration window in the offspring.

At the systemic level, independent of paternal history, we observed that cardiac injury at P1 delays the growth of animals. Similarly, in humans a history of cardiac surgery in infancy is associated with developmental delay and growth problems (Scott & Neal, 2021). However, at the BW level, injured offspring with a paternal history of injury showed increased weight gain and had similar BW to uninjured F1 mice. This increased BW could lead to an increase in blood volume, resulting in the observed eccentric hypertrophy and enhanced cardiac output. In turn, the enhanced cardiac functionality might lead to better organ reperfusion, associated with a reduced EMH demand, and partially to the increase in liver weight (this organ receives up to 25% of the cardiac output and is particularly sensitive to blood flow fluctuations (Møller & Bernardi, 2013). Thus, the improved cardiac functionality in the injured offspring of injured fathers could be the cause of better systemic perfusion, which ameliorates the effects of ischemia on the developing animals. No differences were found at the cardiac or systemic level at later stages, suggesting that the changes observed at weaning were limited to early life. However, we cannot exclude the possibility that long-term consequences might persist in the offspring of injured fathers and affect other biological processes.

Future work will need to address the influence of paternal cardiac injury history on offspring epigenetic changes in cardiac and other cell types under physiological conditions, which may allow a more rapid and robust response to injury. It will also be crucial to understand the genetic or epigenetic mechanism that is altered in the gametes of injured fathers and how this is transmitted throughout the development of the next generation, not only when an injury is performed early in life but also in adulthood.

Approximately 40,000 children require heart surgery each year in the US alone (Pasquali et al., 2020)(Pasquali et al., 2020). Consistent with our mouse model, the finding that neonatal cardiac damage is inherited in mammals may change the way we understand disease and how we conduct anamnesis, as parental cardiac health in early life must be taken into account to understand the health and disease of individuals.

## Author contributions

B.C.: conduction and planning of most of the experiments, supervision of experiments, experimental design, quantification and interpretation of results, conceptualization of the project, writing the manuscript.

M.G.: conduction and planning of experiments.

A.S.M.: conduction of preliminary experiments, experimental design, conceptualization of the project, contribution to the manuscript.

J.L.: conduction of most experimental surgeries.

P.A. snRNA-seq bioinformatics analysis and interpretation, contribution to the manuscript.

L.L.: contribution to snRNA-seq results by providing technical support and protocols, contribution to the manuscript.

E.L.P.: contribution to snRNA-seq results by providing technical support and protocols, contribution to the manuscript.

I.F.: supervised experimental surgeries carried out by J.L., contribution to the manuscript.

N.M.: supervision of all experiments, experimental design, interpretation of results, conceptualization of the project, securing funding, writing the manuscript.

## Material and methods

### KEY RESOURCES TABLE

| REAGENT OR RESOURCE | SOURCE | IDENTIFIER |
| --- | --- | --- |
| **Antibodies** |  |  |
| Troponin T2 | Developmental Studies Hybridoma Bank | Cat# CT3<br>RRID:AB_528495 |
| Myosin, sarcomere (MHC) | Developmental Studies Hybridoma Bank | Cat# MF 20<br>RRID:AB_2147781 |
| Anti-phospho-Histone H3 (Ser10) Antibody, Mitosis Marker | Milipore | Cat#06-570<br>RRID:AB_310177 |
| Wheat Germ Agglutinin 647-Conjugated | Invitrogen | Cat#W32466 |
| Goat anti-Mouse IgG2b Cross-Adsorbed Secondary Antibody, Alexa Fluor™ 488 | Thermo Fisher Scientific | Cat#A-21141<br>RRID:AB_2535778 |
| Goat anti-Mouse IgG Secondary Antibody, Alexa Fluor® 647 conjugate | Thermo Fisher Scientific | Cat# A-21237 |
| Biotin-SP-AffiniPure F(ab')2 Fragment Goat Anti-Rabbit IgG (H+L) | Jackson ImmunoResearch Labs | Cat#111-066-003<br>RRID:AB_2337966 |
| Streptavidin Cy3 conjugate | Thermo Fisher Scientific | Cat#SA1010 |
| Cy™3 Streptavidin | Jackson immunoresearch | Cat# 016-160-084<br>RRID: AB_2337244 |
| Alexa Fluor® 647 anti-mouse TER-119/Erythroid Cells Antibody | BioLegend | Cat# 116218,<br>RRID:AB_528961 |
| **Chemicals, peptides, and recombinant proteins** |  |  |
| DAPI (4',6-Diamidino-2-Phenylindole, Dihydrochloride) | Millipore | Cat#1246530100 |
| Tri reagent | Sigma Aldrich | Cat#T9424-100ML |
| TRI Reagent™ Solution | Invitrogen | Cat#AM9738 |
| 1-bromo-3-chloropropane | Sigma Aldrich | Cat# B9673 |
| Phosphate buffered saline (PBS) | Nzytech | Cat#MB18201 |
| Formalin solution, neutral buffered, 10% | Sigma Aldrich | Cat#HT501128 |
| Sucrose | Sigma Aldrich | Cat#S9378 |
| Isopentane | Merck | Cat#106056 |
| Xylol | Grogg | Cat#G990 |
| p Xylene | Sigma Aldrich | Cat#S8059091 |
| Tween 20 | Sigma Aldrich | Cat#P1379-500ML |
| Triton X-100 | Sigma Aldrich | Cat#T9284-500ML |
| Bovine Serum Albumin (BSA) | Sigma Aldrich | Cat#A7906-100G |
| Goat serum | Milian | Cat#S2000-100 |
| Normal Goat Serum Blocking Solution | Vector Laboratories | Cat# S-1000-20 |
| Citric acid monohydrate (for citrate buffer) | Sigma Aldrich | Cat#C1909-500G |
| Magnesium chloride (MgCl<sub>2</sub>) | Sigma Aldrich | Cat#M8266-100g |
| Spermidine trihydrochloride | Sigma Aldrich | Cat#S2501-1g |
| Tris (Hydroxymethyl) Aminomethane, Free-base, Reagent Grade (500gr) | EMS | Cat#11720 |
| Hydrochloric Acid (HCL) 37% | Grogg | Cat#G840 |
| DTT, Molecular Grade (DL-Dithiothreitol) | Promega | Cat# P1171 |
| Calcium chloride dihydrate (CaCl<sub>2</sub>.2H<sub>2</sub>O) | Sigma Aldrich | Cat#223506-500G |
| Magnesium Acetate | Sigma Aldrich | Cat# M5661 |
| Ethylenediaminetetraacetic acid (EDTA) | Sigma Aldrich | Cat# E6758 |
| Ethylene glycol-bis(2-aminoethylether)-N,N,N',N'-tetraacetic acid EGTA | Sigma Aldrich | Cat# E3889 |
| Trypan blue | Hyclone | Cat# SV30084.01 |
| Direct red80 | Sigma Aldrich | Cat#365548 |
| Picric Acid ((O<sub>2</sub>N)3C<sub>6</sub>H<sub>2</sub>OH) saturated 1,3% | Sigma Aldrich | Cat#P6744-1GA |
| Acetic acid (glacial) 100% | Merk | Cat#1.00063.1000 |
| Ethanol (EtOH) 100% | Grogg | Cat#G003 |
| 2-Propanol | Sigma Aldrich | Cat#67-63-0 |
| Glycerine ≥99.5%, AnalaR NORMAPUR® ACS analytical reagent, redistilled | VWR Chemicals | Cat#56-81-5 |
| **Critical commercial assays** |  |  |
| Vetbond <sup>3M</sup> Tissue adhesive | Vetbond | Cat#1469SB |
| Weigert's Iron Hematoxylin A | EMS | Cat#26044-06 |
| Weigert's Iron Hematoxylin B | EMS | Cat#26044-16 |
| Eosin Y solution | Sigma Aldrich | Cat# 318906 |
| Avidin/Biotin Blocking kit | Vector | Cat#VC-SP-2001-KI01 |
| Fluorescence Mounting Medium | ibidi | Cat#50001_IBI |
| Bouin's Solution (1L) | EMS | Cat#15990-01 |
| Halt Protease Inhibitors Cocktail | ThermoFisher | Cat# 78430 |
| RNase OUT Recombinant Ribonuclease Inhibitor | ThermoFisher | Cat# 10777019 |
| **Experimental models: Organisms/strains** |  |  |
| Mouse: C57BL/6J | The Jackson Laboratory | RRID:IMSR_JAX:000664 |
| **Software and algorithms** |  |  |
| ImageJ | Fiji/ImageJ | RRID:SCR_002285 |
| NDP.view2 Image viewing software | Hamamtsu | Ref#U12388-01 |
| GraphPad Prism 9 | GraphPad Software |  |
| Enrichr | (E. Y. Chen et al., 2013; Kuleshov et al., 2016; Xie et al., 2021) |  |
| ImmGen | (Heng et al., 2008) |  |
| Vevo 2100 Workstation software Version 5.6.1. | Visual Sonics |  |

## Resource availability

### Lead contact

Further information and requests for resources and reagents should be directed to and will be fulfilled by the lead contact, Nadia Mercader.

### Data and code availability

All sequencing data and code will be made available upon peer review.

### Experimental model

All Experiments were approved by the Community of Madrid “Dirección General de Medio Ambiente” in Spain. All animal procedures conformed to EU Directive 86/609/EEC and Recommendation 2007/526/EC regarding the protection of animals used for experimental and other scientific purposes, enforced in Spanish law under Real Decreto 1201/2005. Experiments were conducted under the license PROEX 310_19. Experiments were performed with neonatal and up to one-year-old C57BL/6J mice. Animals were housed together until weaning and then at a density of 4 animals/cage, with the exception of bred males, which were kept isolated after the first breeding. All experiments were performed in a specific pathogen free facility under the following conditions: 20-24°C, 12 hours light-dark cycle and 45-65% relative humidity. Standard chow was provided *ad libitum*. Litter, food, and water were changed regularly.

### Experimental details

Mice were subjected to cardiac injury, sham surgery or cryoanesthetized at P1 (1 day *post-partum*). Injured males were raised to adulthood in parallel with uninjured males from other litters. Upon reaching sexual maturity, male mice were mated with control females. To exclude the possibility that *primiparous* females or a specific combination of breeding could influence the phenotype of the F1 generation, we bred some of the sires and dams more than once and in different combinations. The offspring of uninjured and injured males were then left uninjured or cryoinjured at P1. The F1 generation was sacrificed at different time points depending on the scientific question (7 dpi for CM proliferation; 3 wpp for echocardiography, scar resolution assessment, hypertrophy analysis, and snRNA-seq; and 8-20 wpp for long-term experiments).

## Methods details

### Left ventricular cryoinjury surgery

Prior to surgery, a few drops of surgical glue were applied to the cage near the net to desensitize the mother to its smell. Half of the litter was then separated from the mother and underwent the surgery. Once the pups had woken up and regained their body temperature, they were returned to mother and the other half of the litter was operated on to limit the stress caused to the mother by the separation of the pups. Surgery was performed as described by (Aix et al., 2016). Briefly, P1 mice were anesthetized by hypothermia (*i.e.* cryoanesthesia). A small incision was made in the skin, above the approximate projection of the 3rd and 4th intercostal space. The pectoral muscles were then dissected to expose the thoracic cage and the interior of the chest. With gentle pressure on the abdomen, the apical part of the heart was exposed and approximately 15% of the heart was frozen with a small 24-caliber probe (1 mm) previously immersed in liquid nitrogen. The costal plane was closed, and the skin incision was sutured with surgical glue. Mice were placed on a thermostatic plate to recover body temperature before being returned to the mother. Sham-operated animals underwent the same procedure except for the cryoinjury. Surgery was performed in a blinded fashion.

### Echocardiography

Transthoracic echocardiography was performed in a blinded fashion by an experienced operator using a high-frequency ultrasound system (Vevo 2100, Visualsonics Inc., Canada) with a 30-MHz linear probe. Two-dimensional (2D) and M-mode (MM) echography was performed at a frame rate greater than 230 frames/sec, and pulse wave Doppler (PW) was acquired at a pulse repetition frequency of 40 kHz. Mice were lightly anesthetized with 0.5-2% isoflurane in oxygen, with isoflurane delivery adjusted to maintain a heart rate of 450±50 bpm. Mice were placed in supine position using a heating platform, and heated ultrasound gel was used to maintain normothermia. A basal apex electrocardiogram (ECG) was monitored continuously. Images were transferred to a computer and analyzed off-line using the Vevo 2100 Workstation software. Parasternal standard 2D and MM, long and short axis views (LAX and SAX views, respectively) were acquired to assess left ventricular systolic function. LV ejection fraction, LV fractional shortening, and LV chamber dimensions were calculated from these views.

The mitral valve inflow pattern was assessed using 4-chamber apical PW Doppler echography to evaluate diastolic function. Early and late diastolic velocity peak waves (E and A, respectively), the E/A ratio and isovolumetric relaxation time (IVRT) were measured.

### SnRNA-seq of the whole heart

Whole heart processing was performed according to the protocol of (Cui & Olson, 2020) with some modifications. One whole heart per sample was extracted from euthanized male mice at 21 days post-injury, snap frozen, and stored at −80°C until all samples were available. Two replicates per condition (injured offspring from either uninjured or injured fathers) were used. Cardiac tissue was homogenized and the Ter119 antibody (BioLegend, 1:200) was used to exclude blood cells. After sorting, the integrity of the nuclei was assessed on 5-µl samples diluted 1:2 with trypan blue on the Countess III cell counter (Thermo Fisher). Each nuclear suspension was loaded into a port of a Chromium Next GEM Chip G (10x Genomics) with a target output of 10,000 nuclei. Individual nuclei were encapsulated into emulsion droplets using the Chromium Controller (10x Genomics). SnRNA-seq libraries were prepared using the Chromium Next GEM Single Cell 3’ Kit v3.1 (10x Genomics) and each library was amplified using a SureCycler 8800 thermal cycler (Agilent Technologies). The average size of each library was then calculated using a high sensitivity DNA chip on a 2100 Bioanalyzer (Agilent Technologies) and the concentration was determined using a Qubit fluorometer (Thermo Fisher Scientific). Individual libraries were diluted to 10 nM and pooled for sequencing. The library pool was sequenced at 650 pM in paired-end reads (28 bp Read1, 10 bp Index1, 10 bp Index 2 and 90 bp Read2) on a P3 flow cell (100 cycles) of the NextSeq 2000 platform (Illumina). FastQ files for each sample were generated using the cellranger mkfastq pipeline (10x Genomics). Sequencing was performed in the Genomics Unit of the Spanish National Center for Cardiovascular Research (CNIC).

### snRNA-seq bioinformatics analysis

For bioinformatics analysis we used the *Mus musculus* genome assembly v106 from Ensembl (Cunningham et al., 2022). Cell doublet information was removed using the Scrublet algorithm (Wolock et al., 2019). Counts were then processed using the Seurat pipeline (v4.0) in R (v4.0) (FC et al., 2023; Garnier et al., 2023; Hvitfeldt, 2021; R Core Team, 2018). We removed cells with less than 200 reads per cell. We also removed mitochondrial reads that may have included by contamination. We integrated the samples using the Seurat Feature Integration pipeline and performed clustering of the cells. With the markers obtained after clustering, we assigned the cell types using in-house expertise and public data on cardiac single cell/nuclei RNA-seq (see supplementary methods for the classification). We performed differential expression analysis between control and experimental groups in different cell types with a minimum average of 100 cells using the Libra tool with pseudobulk and EdgeR-LRT-based differential expression (Squair et al., 2021)(. Gene ontology enrichment analysis and pathway analysis were performed using the Clusterprofiler tool (Wu et al., 2021). Graphs were plotted using viridis, palleteer, tidyverse, dittoSand, and UpSetR (Bunis et al., 2020; Garnier et al., 2023; Hvitfeldt, 2021; Wickham et al., 2019; Conway et al., 2017).

### Histological analysis

Organs were immediately dissected after sacrifice, quickly washed in PBS, dried, and measured. They were fixed in 10% formalin for 48 hours at 4°C and then either dehydrated through a graded series of ethanol and embedded in paraffin wax. Paraffin-samples were sectioned at 7 μm using a microtome and mounted on Superfrost Plus slides. Prior to staining, sections were deparaffinized in xylol, and rehydrated through a graded series of ethanol and washed in distilled water.

The following primary and secondary antibodies (Abs) were used for immunofluorescence of paraffin-embedded sections. Primary Abs: mouse anti-Tnnt2 (DSHB, 1:50), mouse anti-Mf20 (Developmental Studies Hybridoma Bank, 1:40), PhH3 (Milipore AB_310177, 1:300). Secondary Abs: anti-WGA 647-conjugated (Invitrogen, 1:200), Alexa Fluor 488, 568, 647 (Thermo Fisher Scientific, 1:250) and biotin anti-rabbit (Jackson Immuno Research, 1:250) followed by incubation with Cy3 streptavidin conjugate (Thermo Fisher Scientific, 1:250). Nuclei were stained with DAPI (Millipore, 1:500) and slides were mounted with Fluorescence Mounting Medium (Ibidi). Sections were rehydrated and antigen retrieved with 100 mM citrate buffer pH 6.0. PBS Triton X-100 0,5% was used for permeabilization and Histoblock (5% BSA, 5% goat serum, 20 mM MgCl_2_ in PBS) to block non-specific binding. Primary Abs were left over night at 4°C, secondary Abs for 1 hour at room temperature. For Ab amplification, endogenous avidin and biotin were blocked with the Avidin/Biotin blocking kit (Vector). All washes were performed with PBS Tween 0.1%.

CM proliferation was quantified as the number of proliferating CMs (PhH3+, Tnnt2/Mf20+) normalized to the area of total nuclei (DAPI+). Eccentric hypertrophy was measured as length of longitudinal CMs in WGA+ Tnnt2+/MF20+ CMs.

Cardiac collagen deposition was assessed by Sirius Red staining. Rehydrated sections were stained in Weigert’s hematoxylin for 8 min, washed in tap water for 10 minutes, stained in Picrosirius red solution (0.1%w/v Direct Red 80 in picric acid) for 1 hour, washed twice in acidified water (0.5% acetic acid), dried, dehydrated with three changes of 100% ethanol, and mounted. Scar resolution was measured as the percentage of collagen area normalized to the cardiac area of the section.

Histological analysis of liver was performed on hematoxylin and eosin-stained sections. After rehydration, sections were stained in Harris haematoxylin for 3 minutes, washed, dipped in 1% acid alcohol solution for a few seconds, rinsed in water, counterstained in 1% eosin for 3-5 minutes, dehydrated in increasing concentrations of alcohol, cleared in xylol, and mounted with mounting medium. Gylcogen deposition and EMH loci were evaluated qualitatively or quantitatively in liver sections.

Images from histological and immunostained sections were acquired with slide scanners (Hamamatsu NanoZoomer 2.0RS and AxioScan Z1) or confocal microscope (Leica SP8 and Zeiss700) using 10x and 20x objectives.

### Quantification and statistical analysis

The statistical analysis tests used in each comparison are shown in the legend of each figure and do not include outlier values. Normal distribution was assessed in each case by the D’Agostino-Pearson normality test prior to selecting the statistical test to be performed, after excluding outliers. GraphPad Prism 9 software was used for statistical tests and plotting. All statistical values are displayed as mean ± standard deviation. All statistics related to snRNA-Seq were performed using the respective packages and core R.

## AKNOWLEDGMENTS

This project has received funding from the European Union’s Horizon 2020 research and innovation programme under grant agreement No 819719 and an Interdisciplinary Grant (UniBe ID Grant) from the University of Bern to N.M. We thank the animal facility, the histology and pathology unit, the advanced imaging unit, and the genomics unit of the Centro Nacional de Investigaciones Cardiovasculares CNIC. The CNIC is supported by the Instituto de Salud Carlos III (ISCIII), the Ministerio de Ciencia e Innovación (MCIN) and the Pro CNIC Foundation and is a Severo Ochoa Center of Excellence (SEV-2015-0505). We are also grateful to MIC-Bern from the University of Bern. B.C. is affiliated to the Graduate School for Cellular and Biomedical Sciences (GCB) of the University of Bern. We thank T. Pedrazzini and V. Corces as well as all the lab member of the N.M. lab for sharing knowledge and advice, and Carlos Garrido Fernández for his help in the setup of the preliminary experiments.

