## Supplemental material and methods for "A paternal cardiac lesion induces cardiac adaptation in offspring"

### Supplemental figures

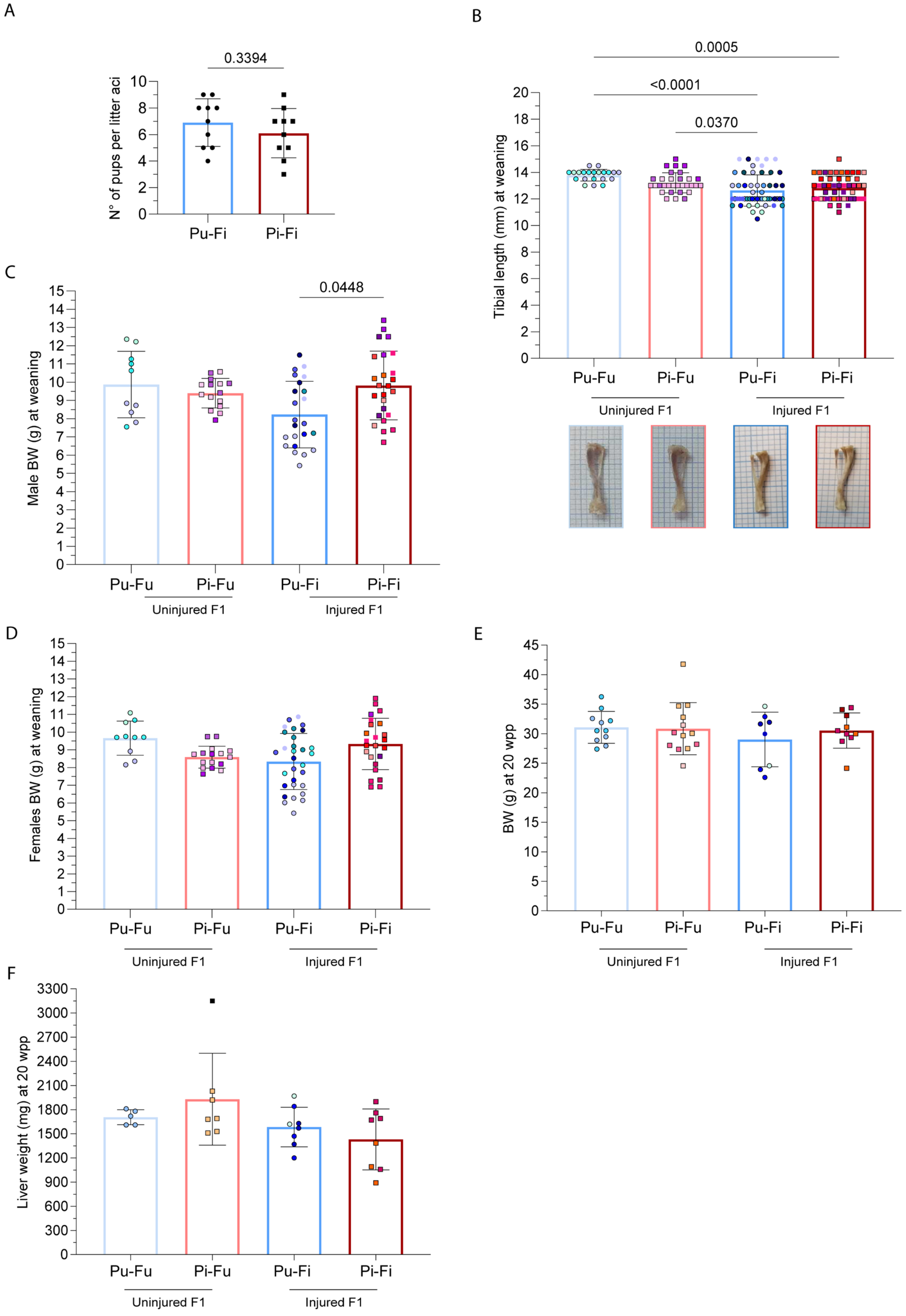


**Figure S1. Systemic changes in injured F1 mice of uninjured and injured fathers at weaning and in adulthood. A**, Number of surviving pups per litter from uninjured and injured fathers after the cryoinjury. Shown is mean and S.D. Unpaired t-test. Aci, after cryoinjury.

**B,** Tibial length (mm) at weaning (3 weeks *post-partum* (wpp)) in uninjured and injured F1 mice. Each bar represents data from siblings of multiple sires, identified by different colors (number of sires: Pu-Fu=3, Pi-Fu=4, Pu-Fi=8, Pi-Fi=8). Shown is mean and S.D. One-way ANOVA, Tukey's multiple comparisons test. On the bottom, representative pictures of tibias from uninjured and injured F1.

**C,** Male F1 BW (g) under physiological condition and after cardiac injury at weaning (3 wpp). Each column represents data from siblings of multiple sires, identified by different colors (number of sires: Pu-Fu=3, Pi-Fu=4, Pu-Fi=5, Pi-Fi=7). Shown is mean and S.D. Kruskal-Wallis test, Dunn's multiple comparisons test.

**D,** Female F1 BW (g) under physiological condition and after cardiac injury at weaning (3 wpp). Each column represents data from siblings of multiple sires, identified by different colors (number of sires: Pu-Fu=3, Pi-Fu=4, Pu-Fi=7, Pi-Fi=8). Shown is mean and S.D. Kruskal-Wallis test, Dunn's multiple comparisons test.

**E,** Body weight of adult F1 mice (20 wpp). Each column represents data from siblings of multiple sires, identified by different colors (number of sires: Pu-Fu=2, Pi-Fu=3, Pu-Fi=3, Pi-FI=3). Shown is mean and S.D. Kruskal-Wallis test, Dunn's multiple comparisons test.

**F,** Liver weight (mg) in adult F1 mice (20 wpp) of uninjured and injured fathers. Each column represents data from siblings of multiple sires, identified by different colors (number of sires: Pu-Fu=1, Pi-Fu=1, Pu-Fi=3, Pi-Fi=2). Shown is mean and S.D. Kruskal-Wallis test, Dunn's multiple comparisons test.

P, paternal generation; F, F1 generation; u, uninjured; I, injured.

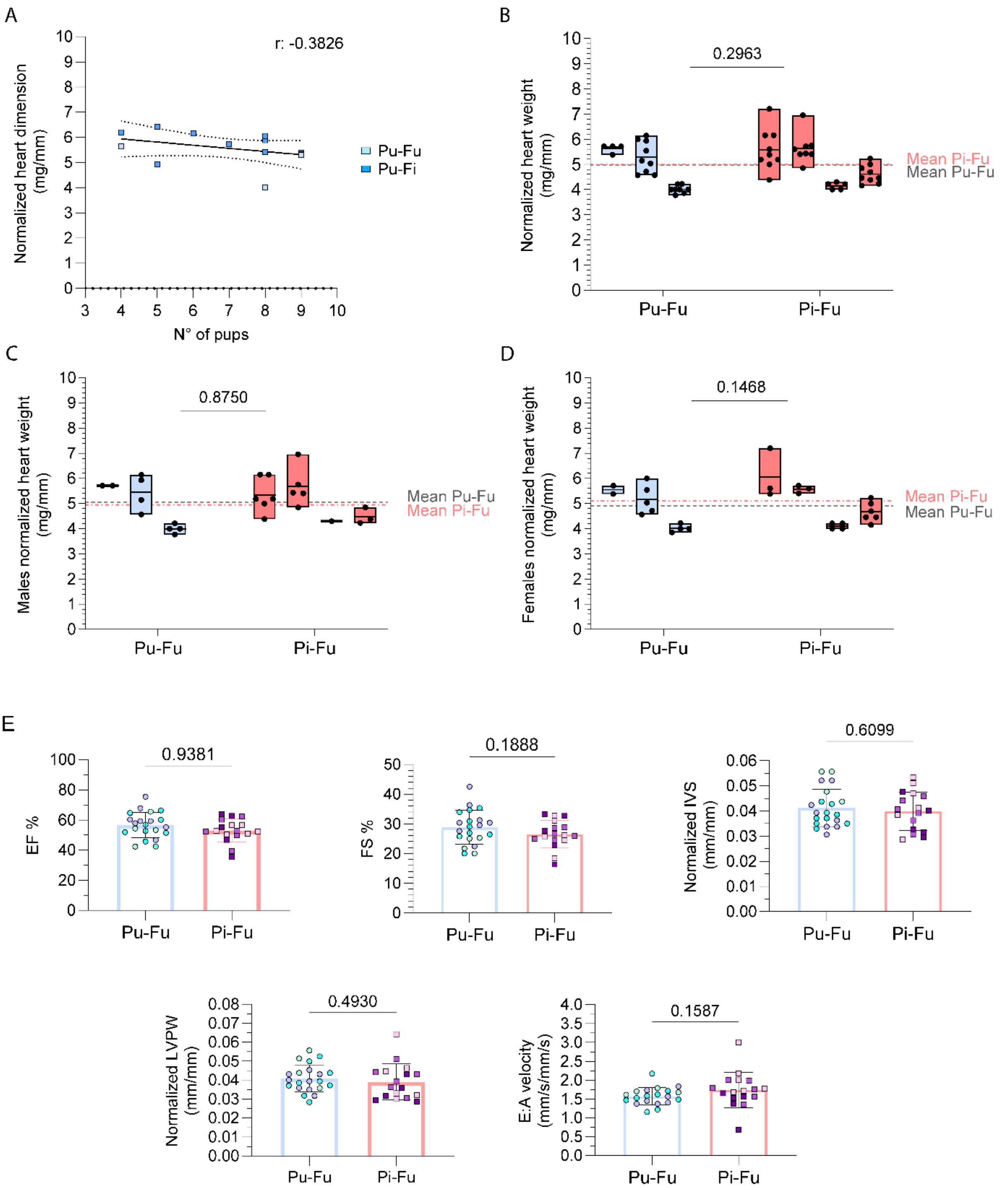


**Figure S2. Cardiac changes in the uninjured offspring of uninjured and injured fathers at weaning.**

**A,** Data on the heart dimension normalized to tibial length (mg/mm) in the control groups (Pu-Fu and Pu-Fi) *vs* number of pups/litter at weaning (3 weeks *post-partum* (wpp). Pearson's correlation test.

**B,** Floating bars representing the normalized heart weight of uninjured F1 mice in different litters (each bar contains siblings of a single family). Two-way ANOVA test.

**C,** Floating bars representing the normalized heart weight of male uninjured F1 mice in different litters (each bar contains siblings of a single family). Two-way ANOVA test.

**D,** Floating bars representing the normalized heart weight of female uninjured F1 mice in different litters (each bar contains siblings of a single family). Two-way ANOVA test.

**E,** Echocardiography of the F1 mice of uninjured and injured fathers under physiological conditions at weaning (3 wpp). Graphs show ejection fraction (EF %), fractional shortening (FS %), interventricular septum (IVS, mm), left ventricle posterior wall (LVPW, mm), and the E (mm/s) A (mm/s) velocity ratio at weaning (3 wpp).

Morphological data are normalized to tibial length. Dots reveal individual hearts from multiple sires, identified by different colors (number of sires: Pu-Fu=3, Pi-Fu=3). Shown is mean and S.D. Unpaired t-test.
P, paternal generation; F, F1 generation; u, uninjured; i, injured.

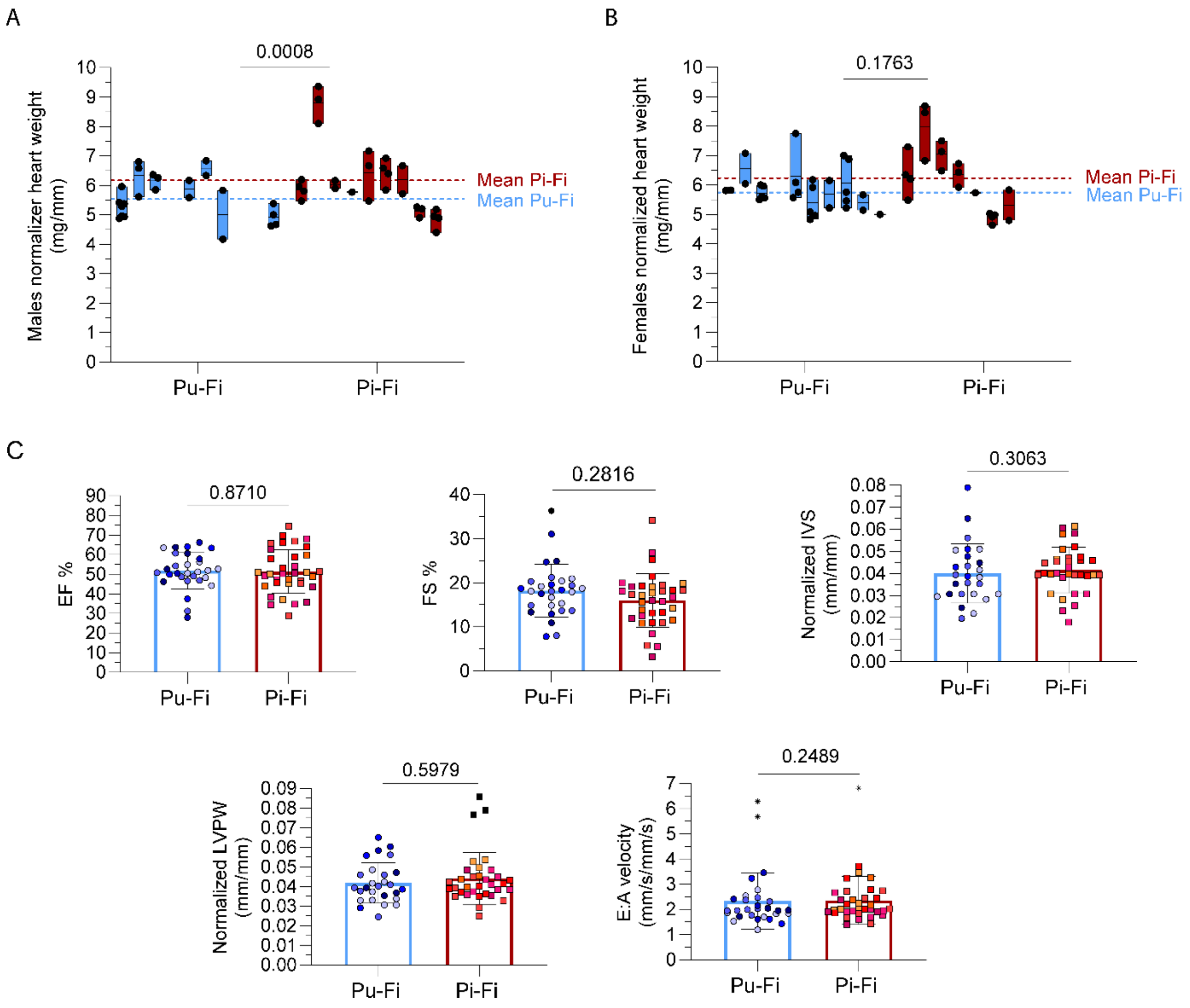


**Supplementary figure 3. Cardiac changes in the injured offspring of uninjured and injured fathers at weaning.**

**A,** Floating bars representing the normalized heart size of male siblings in the different litters (each bar contains siblings of a single family). Dots reveal individual animals. Two-way ANOVA test.

**B,** Floating bars representing the normalized heart size of female siblings in the different litters (each bar contains siblings of a single family). Dots reveal individual animals. Two-way ANOVA test.

**C,** Echocardiography of the injured F1 mice of uninjured and injured fathers at weaning (3 weeks *post-partum*(wpp)). Graphs show ejection fraction (EF %) (unpaired t-test), fractional shortening (FS %) (unpaired t-test), interventricular septum (IVS, mm) (Mann-Whitney test), left ventricle posterior wall (LVPW, mm) (unpaired t-test), and the E (mm/s) A (mm/s) (unpaired t-test) velocity ratio at weaning (3 wpp). Morphological data are normalized to tibial length. Dots reveal individual hearts from multiple sires, identified by different colors (number of sires: Pu-Fi=5, Pi-Fi=6). Shown is mean and S.D.

P, paternal generation; F, F1 generation; u, uninjured; i, injured.

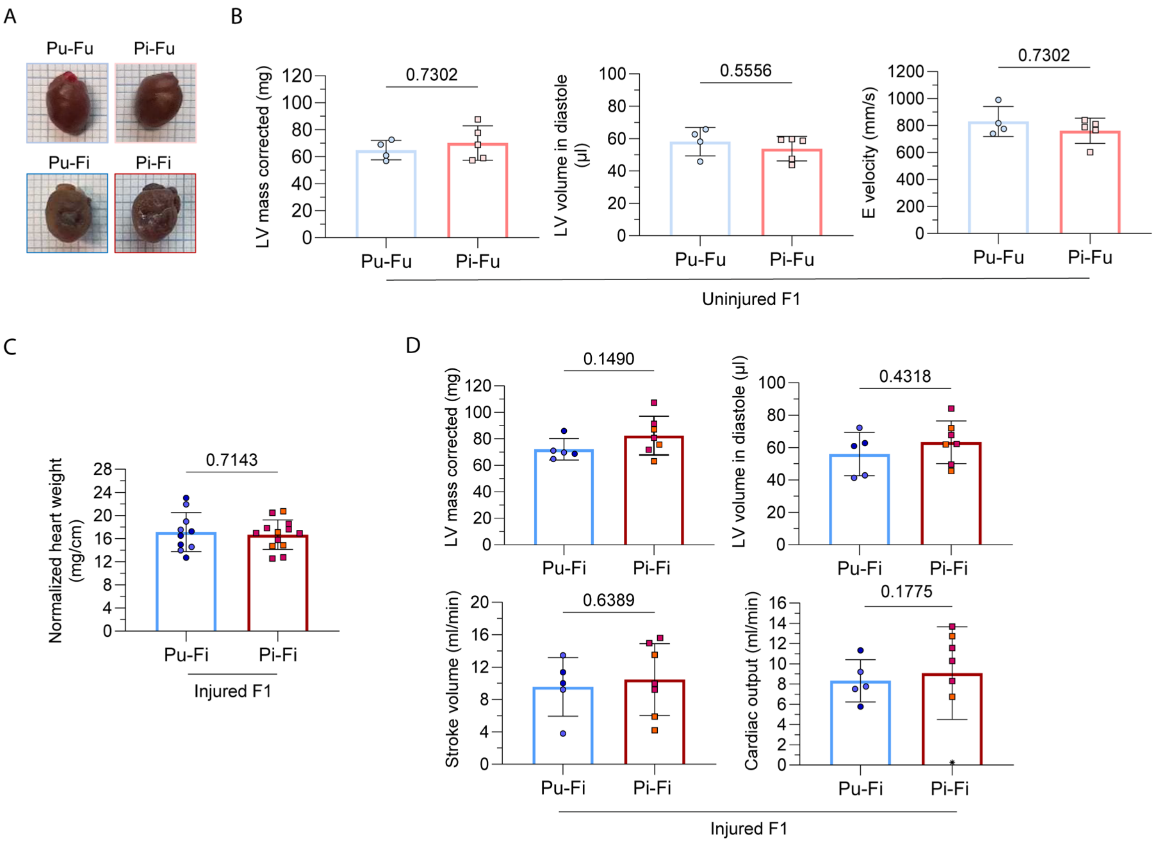


**Figure S4. Cardiac changes in adult offspring of uninjured *vs* injured fathers.**

**A,** Representative hearts of uninjured and injured offspring at 20 wpp.

**B,** Echocardiography assessment of morphological and functional changes in adult uninjured (8 weeks *post-partum* (wpp)) offspring of uninjured and injured fathers. Graphs show LV mass corrected (mg), LV volume in diastole (µl), E velocity (mm/s). Dots show individual hearts from 1 sire per group. Mann-Whitney test.

**C,** Heart weight (g) normalized to body length (cm) of adult (20 wpp) injured F1 mice. Dots show individual hearts from 4 sires (2 Pu-Fi, 2 Pi-Fi). Shown is mean and S.D. Kruskal-Wallis test, Dunn's multiple comparisons test.

**D,** Echocardiography assessment of morphological and functional changes in adult injured (14 wpp) offspring of uninjured and injured fathers. Graphs show left ventricle (LV) mass corrected (mg), LV volume in diastole (µl), stroke volume (ml), cardiac output (ml/min). Dots show individual hearts from 2 sires, one per group. Mann-Whitney test.

P, paternal generation; F, F1 generation; u, uninjured; I, injured; CI, cryoinjury.

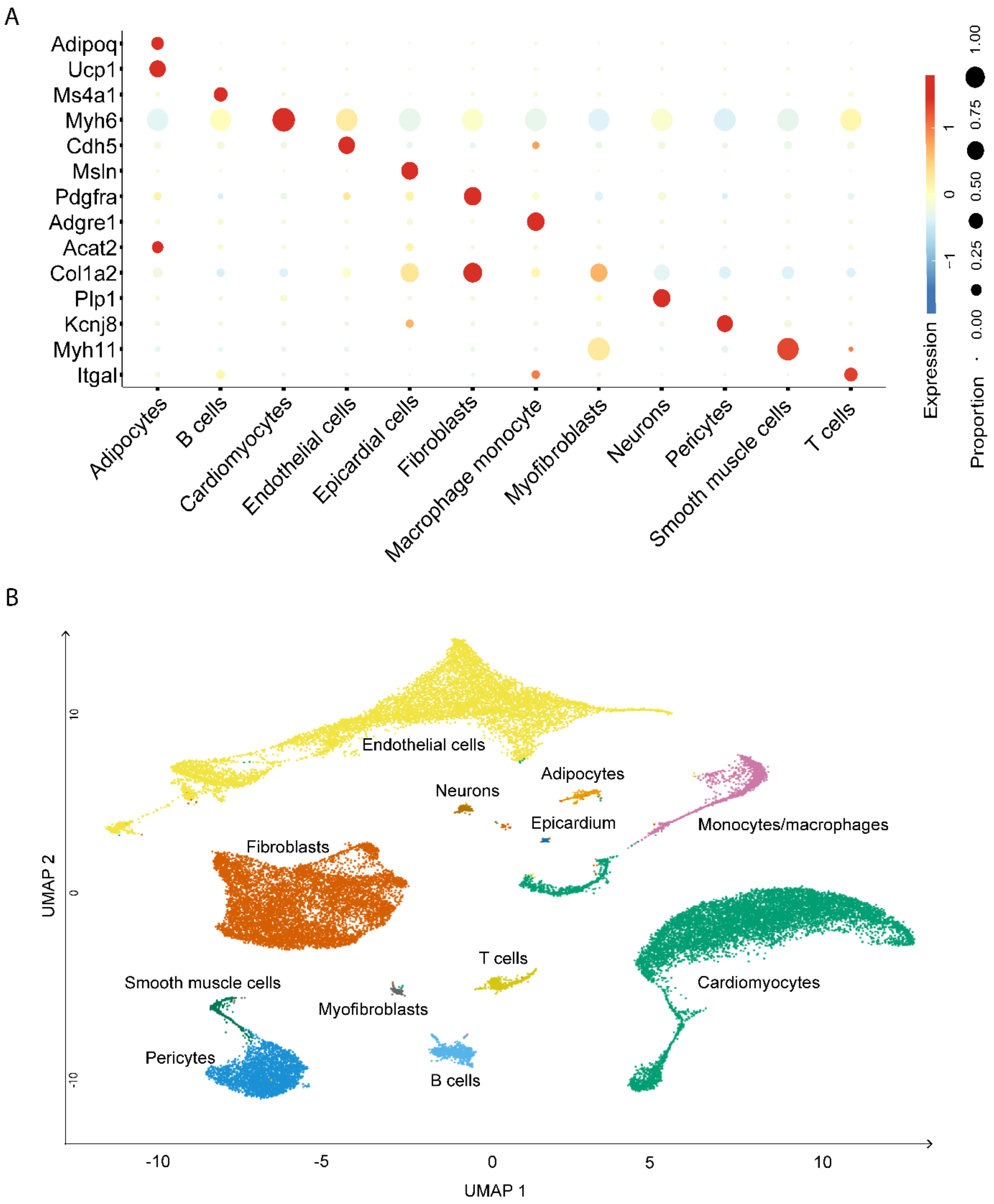


**Figure S5. SnRNA-seq of 21-dpi hearts from uninjured and injured fathers.**
**A,** Bubbleplot of cell population markers used for the identification of snRNA-seq main clusters. Dot color indicates gene expression levels scaled across samples; dot size denotes proportion of cells expressing the gene. **B,** Uniform manifold approximation and projection visualization of unsupervised clustering of 21-dpi whole heart samples from two replicates per condition, displaying subclusters.

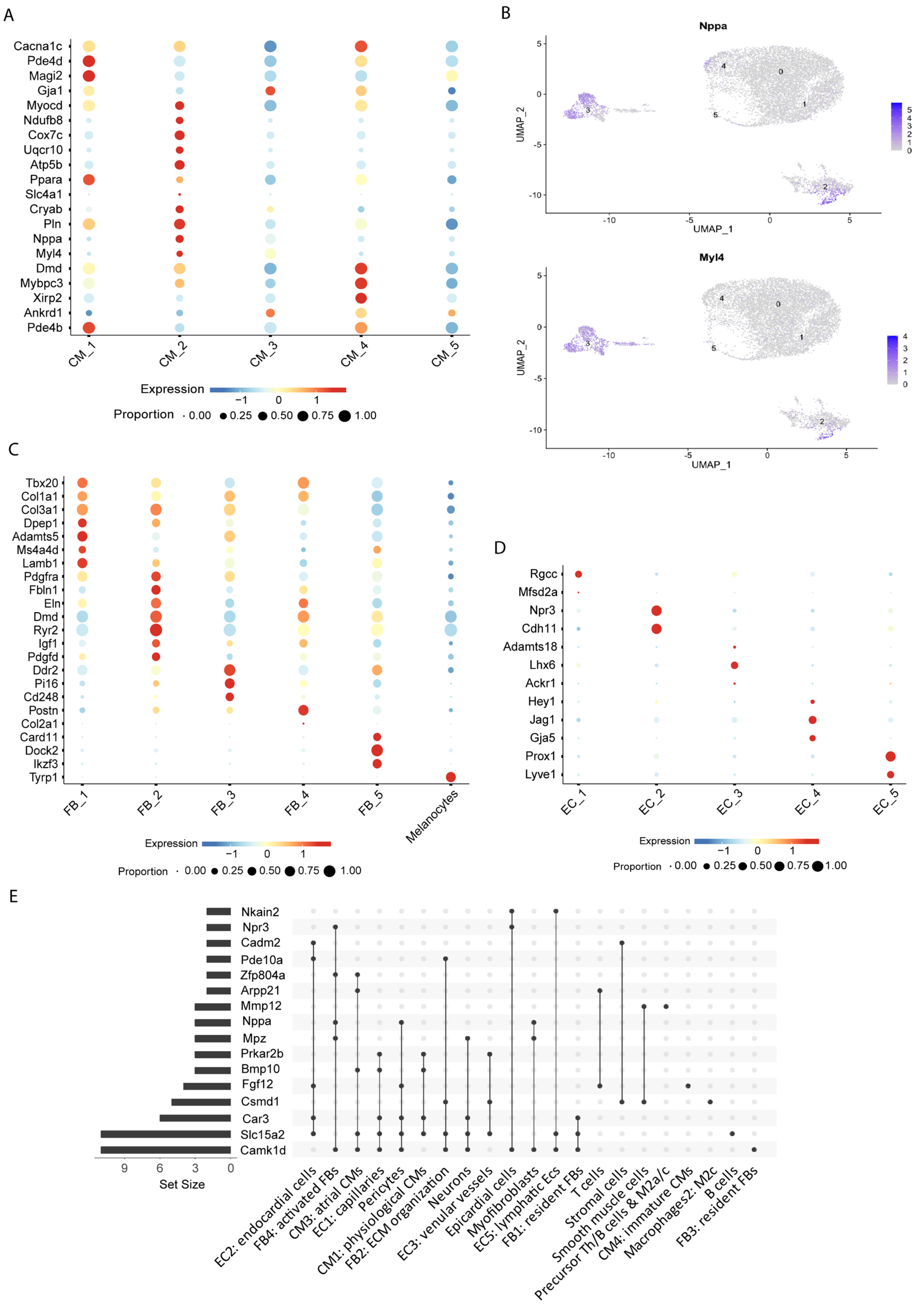


**Figure S6. Identification of snRNA-seq cell populations and differentially expressed genes.**

**A,** Bubbleplot of cell population markers used for the identification of cardiomyocytes (CMs). Dot color indicates gene expression levels scaled across samples; dot size denotes proportion of cells expressing the gene.

**B,** Visualization of the expression of atrial markers *Nppa* and *Myl4* among CM subclusters. Grey to violet scale indicates increasing gene expression levels.

**C,** Bubbleplot of cell population markers used for the identification of fibroblasts (FBs). Dot color indicates gene expression levels scaled across samples; dot size denotes proportion of cells expressing the gene.

**D,** Bubbleplot of cell population markers used for the identification of endothelial cells (ECs). Dot color indicates gene expression levels scaled across samples; dot size denotes proportion of cells expressing the gene.

**E,** UpSet plot of genes found differentially expressed in more than one subpopulation. T cells were plotted as a unique cell group.

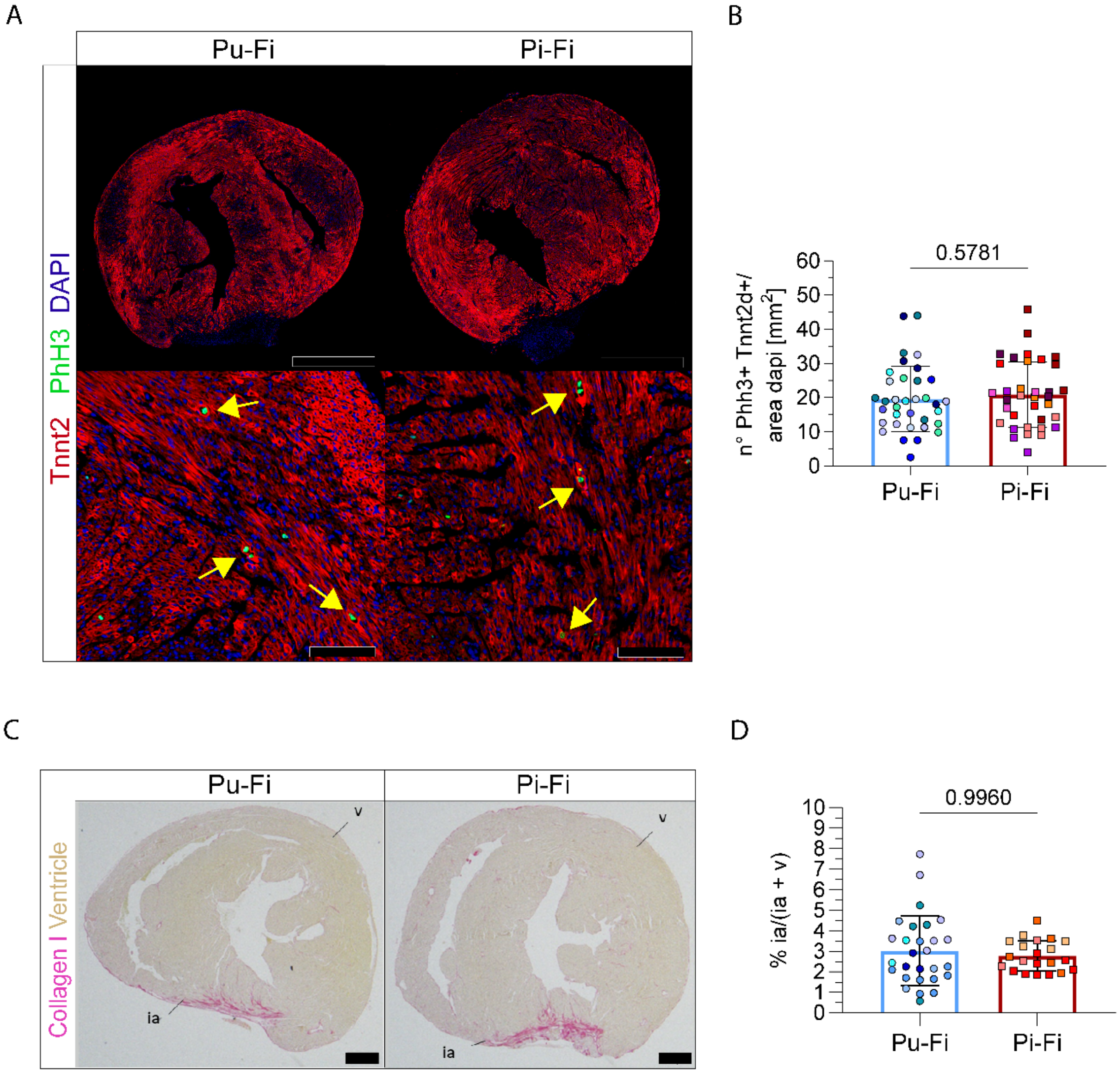


**Figure S7. Cardiac regeneration in the offspring of uninjured *vs* injured fathers.**

**A,** Representative immunofluorescence of F1 hearts at 7 days post-injury (dpi). Tnnt2 marks CMs, PhH3 proliferating cells, DAPI nuclei. Scale bar: 1 mm (top), 100 µm (bottom).

**B,** Quantification of CM proliferation 7 dpi, measured as (PhH3+ Tnnt2+) cells/DAPI area (mm$^2$). Each bar shows data from siblings of multiple sires, identified by different colors (number of sires: Pu-Fi=8, Pi-Fi=7). Shown is mean and S.D. Unpaired t-test.

**C,** Sirius Red stain of hearts of F1 mice at 3 weeks *post-partum*(wpp). Collagen I is stained in pink and identifies the injury area (ia), the rest of the ventricle (v) is stained in clear brown. Scale bar: 100 µm.

**D,** Quantification of scar resolution at 3 wpp measured as the percentage of injured area (ia) to the whole ventricles area (ia+ v). Each bar shows data from siblings of multiple sires, identified by different colors (number of sires: Pu-Fi=6, Pi-Fi=4). Shown is mean and S.D. Mann-Whitney test. P, paternal generation; F, F1 generation; u, uninjured; i, injured; ia, injury area; v, ventricle.

**Supplemental tables**

| Cell_type | Gene | Avg_logFC | P_val | P_val_adj |
|---|---|---|---|---|
| Adipocytes | Ryr3 | -6.28608 | 5.75E-08 | 7.96E-04 |
| B_cells | Slc15a2 | 7.26579 | 6.66E-09 | 9.40E-05 |
| B_cells | Dpp10 | 4.57657 | 4.37E-07 | 3.08E-03 |
| CM_1 | Slc15a2 | 5.62961 | 9.80E-28 | 1.71E-23 |
| CM_1 | Bmp10 | 4.84569 | 1.84E-07 | 5.36E-04 |
| CM_1 | Car3 | 3.96076 | 3.08E-12 | 2.69E-08 |
| CM_1 | Tspan2 | 2.34268 | 5.04E-09 | 2.20E-05 |
| CM_1 | Prkar2b | 2.01485 | 1.14E-07 | 3.97E-04 |
| CM_1 | Gck | -1.26597 | 6.81E-06 | 1.49E-02 |
| CM_1 | Ucp3 | 1.01071 | 2.01E-05 | 3.51E-02 |
| CM_3 | Arpp21 | 6.63361 | 2.16E-06 | 2.71E-03 |
| CM_3 | Ppfia2 | 6.17054 | 4.86E-05 | 3.17E-02 |
| CM_3 | Zfp804a | -4.80845 | 3.17E-07 | 5.95E-04 |
| CM_3 | Slc15a2 | 4.40883 | 5.12E-07 | 8.55E-04 |
| CM_3 | Ms4a4b | -4.15908 | 6.25E-05 | 3.24E-02 |
| CM_3 | Eya1 | 2.95929 | 2.15E-07 | 4.61E-04 |
| CM_3 | Adm | 2.91797 | 3.55E-05 | 2.67E-02 |
| CM_3 | Bmp10 | 2.51627 | 3.25E-11 | 2.44E-07 |
| CM_3 | Stk32a | 2.47425 | 1.12E-08 | 5.61E-05 |
| CM_3 | Cdcp3 | 2.33982 | 1.34E-11 | 2.01E-07 |
| CM_3 | Eva1c | 2.12877 | 5.70E-05 | 3.17E-02 |
| CM_3 | Tgfb1i1 | -1.83522 | 1.98E-05 | 1.81E-02 |
| CM_3 | Myl9 | -1.80003 | 1.04E-06 | 1.42E-03 |
| CM_3 | Vdr | 1.78084 | 4.50E-05 | 3.07E-02 |
| CM_3 | Rfx2 | 1.71447 | 2.98E-08 | 1.12E-04 |
| CM_3 | Txnip | 1.61291 | 8.19E-05 | 3.78E-02 |
| CM_3 | Pdlim7 | -1.57862 | 1.27E-05 | 1.36E-02 |
| CM_3 | Arhgef26 | 1.56619 | 3.88E-05 | 2.77E-02 |
| CM_3 | Shroom4 | 1.50079 | 5.63E-05 | 3.17E-02 |
| CM_3 | Nmrk2 | -1.45818 | 8.32E-05 | 3.78E-02 |
| CM_3 | Vwc2 | -1.43308 | 1.10E-07 | 2.75E-04 |
| CM_3 | Mgmt | 1.43303 | 9.03E-05 | 3.87E-02 |
| CM_3 | Mbp | 1.41240 | 5.77E-07 | 8.67E-04 |
| CM_3 | Il31ra | 1.38581 | 1.09E-04 | 4.54E-02 |
| CM_3 | Kirrel3 | -1.37746 | 5.38E-08 | 1.62E-04 |
| CM_3 | Kctd1 | 1.14155 | 2.05E-05 | 1.81E-02 |
| CM_3 | Plxdc2 | -1.13367 | 2.27E-05 | 1.89E-02 |
| CM_3 | Kazn | 1.09943 | 7.23E-05 | 3.50E-02 |
| CM_3 | Ror2 | 1.09895 | 6.03E-05 | 3.23E-02 |
| CM_3 | Thsd7a | -1.06086 | 3.37E-05 | 2.66E-02 |
| CM_3 | Camk1d | 1.05679 | 2.85E-06 | 3.29E-03 |
| CM_3 | Daam2 | 1.03096 | 1.47E-05 | 1.48E-02 |
| CM_3 | Hexb | 1.02612 | 5.16E-05 | 3.17E-02 |
| CM_3 | Sh3bgr | -1.00847 | 5.59E-05 | 3.17E-02 |

| | | | | |
|---|---|---|---|---|
| CM_4 | Shisa6 | 6.90661 | 1.46E-07 | 1.09E-03 |
| CM_4 | Fgf12 | 1.80730 | 7.37E-07 | 3.67E-03 |
| CM_5 | Adgrb3 | 6.91149 | 1.32E-07 | 1.62E-03 |
| EC_1 | Cidec | 6.39998 | 2.55E-06 | 6.15E-03 |
| EC_1 | Ankef1 | 6.19355 | 5.01E-06 | 1.06E-02 |
| EC_1 | Ptger3 | 6.12902 | 8.55E-06 | 1.45E-02 |
| EC_1 | Slc15a2 | 5.62384 | 1.63E-19 | 2.75E-15 |
| EC_1 | Bmp10 | 4.45637 | 2.75E-07 | 9.27E-04 |
| EC_1 | Car3 | 4.31169 | 1.37E-16 | 1.16E-12 |
| EC_1 | Ntrk2 | 3.55633 | 8.57E-06 | 1.45E-02 |
| EC_1 | Cfd | 3.25684 | 2.84E-05 | 3.68E-02 |
| EC_1 | Prkar2b | 2.01111 | 1.70E-07 | 7.16E-04 |
| EC_1 | Camk1d | 1.04675 | 7.04E-11 | 3.96E-07 |
| EC_2 | Car3 | 6.22989 | 1.66E-05 | 1.45E-02 |
| EC_2 | Slc15a2 | 4.73889 | 3.24E-08 | 1.02E-04 |
| EC_2 | Cadm2 | -2.51742 | 2.35E-06 | 3.39E-03 |
| EC_2 | Dlgap2 | -2.48585 | 1.83E-05 | 1.45E-02 |
| EC_2 | Rtl4 | -2.17033 | 5.00E-06 | 6.09E-03 |
| EC_2 | Cdh4 | 1.85009 | 1.41E-08 | 5.58E-05 |
| EC_2 | Ppp2r2b | -1.74411 | 3.84E-12 | 3.04E-08 |
| EC_2 | Nmnat2 | 1.60016 | 2.87E-06 | 3.78E-03 |
| EC_2 | Cmah | 1.58263 | 2.84E-07 | 5.63E-04 |
| EC_2 | Adamtsl2 | -1.44581 | 5.97E-07 | 1.05E-03 |
| EC_2 | Fgf12 | 1.42115 | 7.22E-06 | 8.17E-03 |
| EC_2 | Col25a1 | 1.39016 | 1.37E-14 | 2.17E-10 |
| EC_2 | Ptprt | -1.38794 | 2.67E-07 | 5.63E-04 |
| EC_2 | Pxylp1 | -1.34648 | 2.97E-05 | 2.24E-02 |
| EC_2 | Fam107b | 1.24272 | 4.27E-05 | 3.07E-02 |
| EC_2 | Opcml | -1.23520 | 4.58E-05 | 3.15E-02 |
| EC_2 | Pde10a | 1.21788 | 3.98E-10 | 2.10E-06 |
| EC_2 | Errfi1 | -1.09837 | 1.74E-05 | 1.45E-02 |
| EC_3 | Prkar2b | 4.84856 | 9.45E-07 | 4.61E-03 |
| EC_3 | Slc15a2 | 4.69549 | 2.82E-07 | 2.18E-03 |
| EC_3 | Ranbp1 | -3.13696 | 1.12E-05 | 2.50E-02 |
| EC_3 | Ntrk3 | 2.81437 | 4.61E-06 | 1.35E-02 |
| EC_3 | Csmd1 | 2.79540 | 4.00E-06 | 1.35E-02 |
| EC_3 | 2610002M06Rik | -2.55573 | 1.20E-05 | 2.50E-02 |
| EC_3 | Robo2 | -1.00116 | 2.18E-05 | 3.99E-02 |
| EC_4 | Itga8 | -6.08333 | 4.20E-06 | 2.79E-02 |
| EC_4 | Chrm3 | -1.72751 | 1.31E-07 | 1.74E-03 |
| EC_5 | Slc15a2 | 6.26860 | 1.11E-05 | 1.98E-02 |
| EC_5 | Ckap2 | 4.22568 | 1.01E-05 | 1.98E-02 |
| EC_5 | Gria2 | -3.80526 | 6.77E-06 | 1.93E-02 |
| EC_5 | Nkain2 | -2.64043 | 1.14E-08 | 1.63E-04 |
| EC_5 | Stab2 | 2.46440 | 9.26E-06 | 1.98E-02 |
| EC_5 | F8 | 2.22802 | 5.15E-06 | 1.84E-02 |

| | | | | |
|---|---|---|---|---|
| EC_5 | Cenpf | 2.18822 | 2.88E-05 | 4.12E-02 |
| EC_5 | Macrod2 | 1.26760 | 2.14E-05 | 3.39E-02 |
| EC_5 | Camk1d | 1.21733 | 3.31E-08 | 2.36E-04 |
| EC_5 | Lrmda | 1.19371 | 5.13E-06 | 1.84E-02 |
| Epicardium | Nkain2 | 7.65591 | 6.96E-06 | 1.92E-02 |
| Epicardium | Grin2a | 7.54881 | 1.41E-05 | 1.92E-02 |
| Epicardium | Tmeff2 | 7.53967 | 1.46E-05 | 1.92E-02 |
| Epicardium | Bcl2 | 7.50415 | 1.82E-05 | 2.08E-02 |
| Epicardium | Zfp804b | 7.36232 | 4.03E-05 | 2.98E-02 |
| Epicardium | Adamts9 | 4.09114 | 8.01E-06 | 1.92E-02 |
| Epicardium | Bach2 | 3.89064 | 3.22E-05 | 2.89E-02 |
| Epicardium | Stim1 | 3.82663 | 4.98E-05 | 3.47E-02 |
| Epicardium | Slc25a21 | -3.69490 | 3.62E-05 | 2.90E-02 |
| Epicardium | Kcnma1 | -3.44723 | 5.38E-07 | 3.38E-03 |
| Epicardium | Osbpl6 | -2.80560 | 5.43E-05 | 3.59E-02 |
| Epicardium | Hpgd | 2.80136 | 1.24E-05 | 1.92E-02 |
| Epicardium | Ccnd3 | 2.77640 | 1.53E-05 | 1.92E-02 |
| Epicardium | Clstn2 | 2.53689 | 1.15E-05 | 1.92E-02 |
| Epicardium | Kiz | 2.46032 | 2.19E-05 | 2.29E-02 |
| Epicardium | Npr3 | 2.44414 | 6.40E-05 | 4.02E-02 |
| Epicardium | Ebf1 | 2.42440 | 7.36E-05 | 4.40E-02 |
| Epicardium | Magi3 | 1.99223 | 3.69E-05 | 2.90E-02 |
| Epicardium | Gpc3 | 1.96673 | 9.05E-06 | 1.92E-02 |
| Epicardium | Camk1d | 1.89051 | 3.41E-07 | 3.38E-03 |
| Epicardium | Lhfp | 1.87511 | 2.70E-05 | 2.61E-02 |
| FB_1 | Slc15a2 | 4.81101 | 1.72E-13 | 2.85E-09 |
| FB_1 | Car3 | 4.55628 | 6.25E-11 | 3.45E-07 |
| FB_1 | Cidea | 3.25110 | 1.46E-05 | 4.02E-02 |
| FB_1 | Camk1d | 1.19922 | 1.10E-12 | 9.10E-09 |
| FB_1 | Acta2 | -1.07284 | 1.02E-05 | 3.48E-02 |
| FB_2 | Slc15a2 | 7.06457 | 1.68E-08 | 1.39E-04 |
| FB_2 | Csmd1 | 3.73199 | 1.17E-06 | 4.83E-03 |
| FB_2 | Junb | 2.38446 | 2.09E-06 | 4.93E-03 |
| FB_2 | Sorcs1 | 1.86617 | 6.55E-06 | 1.20E-02 |
| FB_2 | Acer2 | 1.32780 | 2.31E-05 | 3.48E-02 |
| FB_2 | Hunk | -1.22760 | 7.56E-06 | 1.25E-02 |
| FB_2 | Grip1 | 1.22447 | 3.52E-05 | 4.69E-02 |
| FB_2 | Camk1d | 1.17932 | 3.25E-11 | 5.37E-07 |
| FB_2 | Ptpn13 | 1.07941 | 3.32E-06 | 6.87E-03 |
| FB_2 | Pde10a | 1.04185 | 2.00E-06 | 4.93E-03 |
| FB_2 | Kcnc2 | 1.02126 | 1.51E-06 | 4.93E-03 |
| FB_3 | Il1rapl1 | -1.88251 | 6.93E-06 | 3.45E-02 |
| FB_3 | Gfra1 | 1.62706 | 2.99E-11 | 2.23E-07 |
| FB_3 | Camk1d | 1.11433 | 2.62E-12 | 3.92E-08 |
| FB_4 | Mpz | 6.21210 | 4.69E-06 | 1.01E-02 |
| FB_4 | Zfp804a | 6.19851 | 1.20E-05 | 2.00E-02 |

| | | | | |
|---|---|---|---|---|
| FB_4 | Acss3 | 3.67781 | 2.66E-06 | 7.97E-03 |
| FB_4 | Tenm2 | 3.33536 | 4.59E-06 | 1.01E-02 |
| FB_4 | Tll1 | 2.95585 | 7.70E-07 | 2.89E-03 |
| FB_4 | Nrxn3 | 2.65597 | 1.70E-09 | 8.50E-06 |
| FB_4 | Npr3 | 2.06996 | 1.19E-09 | 8.50E-06 |
| FB_4 | Nppa | 1.74457 | 2.01E-05 | 2.83E-02 |
| FB_4 | Prkch | 1.53293 | 8.77E-06 | 1.64E-02 |
| FB_4 | Camk1d | 1.14384 | 4.46E-10 | 6.69E-06 |
| Macrophages_1 | Spp1 | -7.60556 | 1.17E-14 | 5.47E-11 |
| Macrophages_1 | Gpnmb | -6.56864 | 7.26E-24 | 5.08E-20 |
| Macrophages_1 | Elovl6 | 6.29868 | 1.07E-05 | 1.15E-02 |
| Macrophages_1 | Atp6v0d2 | -5.09654 | 1.02E-09 | 3.59E-06 |
| Macrophages_1 | Mamdc2 | -4.28752 | 6.30E-25 | 8.82E-21 |
| Macrophages_1 | Kctd8 | -3.85247 | 2.11E-06 | 3.28E-03 |
| Macrophages_1 | Vsig4 | 3.40921 | 6.67E-06 | 8.49E-03 |
| Macrophages_1 | Mmp12 | -3.03740 | 1.66E-05 | 1.49E-02 |
| Macrophages_1 | St18 | -2.98488 | 2.67E-08 | 7.28E-05 |
| Macrophages_1 | Adgb | 2.76619 | 8.78E-06 | 1.03E-02 |
| Macrophages_1 | Fam20c | -2.32193 | 1.95E-06 | 3.28E-03 |
| Macrophages_1 | Kcnip4 | 1.96050 | 1.12E-06 | 2.24E-03 |
| Macrophages_1 | Klra2 | 1.76559 | 5.34E-05 | 4.40E-02 |
| Macrophages_1 | Fat3 | -1.72797 | 3.12E-08 | 7.28E-05 |
| Macrophages_1 | St3gal5 | 1.43755 | 1.70E-05 | 1.49E-02 |
| Macrophages_1 | Adgrl3 | 1.32167 | 1.16E-05 | 1.16E-02 |
| Macrophages_1 | Myo1e | -1.23221 | 3.96E-06 | 5.55E-03 |
| Macrophages_2 | Csmd1 | 2.81631 | 1.07E-05 | 4.86E-02 |
| Macrophages_2 | Gda | 2.29316 | 1.00E-06 | 6.82E-03 |
| Macrophages_2 | Crebrf | 1.98674 | 2.86E-08 | 3.90E-04 |
| Monocytes | Col4a6 | 3.42806 | 4.21E-06 | 4.58E-02 |
| Neurons | Slc15a2 | 6.76578 | 5.67E-08 | 3.72E-04 |
| Neurons | Car3 | 6.61893 | 1.90E-06 | 8.33E-03 |
| Neurons | Mpz | 4.22148 | 4.65E-13 | 6.11E-09 |
| Neurons | Rad51b | 3.21354 | 3.41E-06 | 1.12E-02 |
| Neurons | Camk1d | 1.23594 | 4.74E-06 | 1.24E-02 |
| Pericyte | Slc15a2 | 5.10428 | 4.23E-24 | 6.70E-20 |
| Pericyte | Car3 | 3.13462 | 1.25E-05 | 2.83E-02 |
| Pericyte | Thrsp | 3.11036 | 1.06E-05 | 2.81E-02 |
| Pericyte | Nppa | 1.28605 | 4.10E-07 | 1.62E-03 |
| Pericyte | Camk1d | 1.10268 | 4.28E-12 | 3.39E-08 |
| Pericyte | Pparg | 1.07778 | 4.71E-08 | 2.49E-04 |
| Pericyte | Fgf12 | 1.03115 | 1.69E-06 | 5.36E-03 |
| Precursor_Th_b | Slc9a7 | 6.93916 | 3.67E-07 | 2.11E-03 |
| Precursor_Th_b | Mmp12 | -6.93440 | 4.90E-08 | 5.62E-04 |
| Smooth_muscle_cells | Csmd1 | 3.25169 | 1.07E-09 | 1.43E-05 |
| Smooth_muscle_cells | Camk1d | 1.27241 | 2.91E-07 | 1.94E-03 |
| Myofibroblasts | Mpz | 6.87103 | 1.04E-05 | 2.20E-02 |

| | | | | |
|---|---|---|---|---|
| Myofibroblasts | Gria4 | 4.01041 | 1.05E-05 | 2.20E-02 |
| Myofibroblasts | Slit2 | 3.08436 | 1.50E-08 | 1.89E-04 |
| Myofibroblasts | Nppa | 2.97953 | 7.56E-07 | 4.74E-03 |
| Myofibroblasts | Negr1 | 1.90778 | 1.24E-05 | 2.22E-02 |
| Myofibroblasts | Eln | -1.43351 | 2.14E-06 | 8.95E-03 |
| Myofibroblasts | Camk1d | 1.15351 | 7.49E-06 | 2.20E-02 |
| Stromal_1 | Csmd1 | 8.31949 | 1.20E-17 | 1.46E-13 |
| Stromal_1 | Cdh19 | 6.66612 | 7.04E-08 | 2.86E-04 |
| Stromal_1 | Sv2b | -6.49585 | 4.09E-07 | 8.30E-04 |
| Stromal_1 | Scn7a | 6.33094 | 1.60E-06 | 2.78E-03 |
| Stromal_1 | Cadm2 | 4.61667 | 1.62E-07 | 4.93E-04 |
| Stromal_1 | Lsamp | -4.27923 | 3.43E-06 | 4.64E-03 |
| Stromal_1 | Nrg1 | 4.21605 | 1.01E-10 | 6.16E-07 |
| Stromal_1 | Cdh11 | 4.10242 | 2.20E-05 | 2.23E-02 |
| Stromal_1 | Ddhd1 | 3.14131 | 1.84E-05 | 2.03E-02 |
| Stromal_1 | Slc35f1 | 2.81873 | 2.75E-06 | 4.18E-03 |
| Stromal_1 | Rasgef1b | 2.66404 | 2.48E-07 | 6.03E-04 |
| Stromal_1 | Pde3a | 2.47067 | 6.55E-06 | 7.98E-03 |
| Stromal_2 | Col8a1 | -3.66016 | 8.49E-12 | 1.05E-07 |
| T_cells | Arpp21 | 8.13970 | 4.20E-10 | 4.23E-06 |
| T_cells | C7 | -4.83610 | 1.77E-08 | 8.21E-05 |
| T_cells | Ncr1 | -3.36398 | 9.28E-07 | 3.22E-03 |
| T_cells | Fgf12 | 2.90248 | 6.09E-10 | 4.23E-06 |

**Table S1: list of differentially expressed genes within subclusters.**
SnRNA-seq list of differentially expressed genes (DEG) identified by EdgeR after pseudobulk. DEG with LogFC <-1 or >+1 and p adjusted ≤ 0.05 were identified within each subcluster. Highlighted in red genes commonly found differentially expressed in more than one cluster.

**Supplemental methods**

The main clusters classification identified CMs (*Myh6*), followed by ECs (*Cdh5*), FBs (*Pdgfra* and *Col1a2*), pericytes (*Kcnj8*), myofibroblasts (*Acta2*, *Col1a2*), smooth muscle cells (*Myh11*), epicardial cells (*Msln*), neurons (*Plp1*), adipocytes (*Adipoq, Ucp1*), and cells of the immune system, including B cells (*Ms4a1*), T cells (*Itgal*) and monocytes/macrophages (*Adgre1* and *Itgal*).

Subclustering analysis allowed the identification of subpopulation within the main clusters (Figure S5). CMs subcluster 1 was identified as the main and most abundant CM cell type, expressing genes involved in heart contraction (*Cacna1c*, *Pde4d*), cell junction assembly (*Magi2*, *Gja1*), and muscle system process (*Pde4d*, *Myocd*). CMs subcluster 2 expressed genes involved in aerobic respiration, wit h activation of OXPHOS (*Ndufb8*, *Cox7c*, *Uqcr10*, *Atp5b*) and ATP metabolic processes (*Ppara*, *Slc4a1*). In addition, by checking the expression of *Cryab* and *Pln,* genes upregulated in response to high workload (Litviňuková et al., 2020a), they were also specifically higher in CMs subcluster 2. CMs subcluster 3 was assigned to atrial CMs based on the increased expression of *Nppa* and *Myl4* cell markers (Tucker et al., 2020). CMs subcluster 4 was a smaller population expressing genes involved in heart development and differentiation (*Sorpbs2*, *Dmd*, *Mybpc3*, *Ttn*), together with genes induced upon injury (*Xirp2*, *Ankrd1*, *Pde4b*) (Cui et al., 2020; S. Li et al., 2022). This genetic profile suggests that CMs subcluster 4 could be an immature CMs population, possibly emerging in response to the injury and differentiating into functional CMs to regenerate the damaged tissue. CMs subcluster 5 was the smallest CMs subpopulation, composed of approximately 1% of all CMs, to which no particular function could be attributed based on the gene signature.

ECs classification was based on previously described markers, and allowed us to identify capillaries (subcluster 1; *Rgcc* and *Mfsd2a;* (Kalucka et al., 2020; Vanlandewijck et al., 2018), endocardial cells (subcluster 2; *Npr3*, *Cdh11;* (Feng et al., 2019; Y. Li et al., 2018; McCracken et al., 2022), venular vessels (subcluster 3, *Adamts18*, *Lhx6*, *Ackr1;* He et al., 2022; Trimm & Red-Horse, 2023), arteriole ECs (subcluster 4; *Hey1*, *Jag1*, *Gja5*, (He et al., 2022; McCracken et al., 2022)), and lymphatic ECs (subcluster 5; *Prox1*, *Lyve1;* (Kalucka et al., 2020; Lee, 2020)).

Within FB subclusters, we could define two sub-populations of resident FBs: FBs subcluster 1 and 3. FBs subcluster 1 expressed previously described canonical fibroblast genes, such as *Tbx20*, *Col1a1*, *Col3a1*, *Dpep1*, *Adamts5*, *Ms4a4d*, *Lamb1* (Skelly et al., 2018). FBs subcluster 2 expressed genes involved in extracellular matrix (ECM) organization and muscle tissue process/development, such as *Pdgfra*, *Col3a1*, *Fbln1*, *Eln*, *Dmd*, *Ryr2*, *Igf1*, *Pdgfd*, suggesting a possible role in heart healing processes. FBs subcluster 3 was characterized by the expression of *Ddr2*, *Pi16*, *Cd248*, and *Efhd1*, previously described to characterize FBs involved in physiological function of the heart and other organs (Farbehi et al., 2019; Lendahl et al., 2022; Z. Wang et al., 2020). FBs subcluster 4 corresponded to an activated FB state, expressing high *Postn*, and *Col1a1, Col2a1* genes. FBs subcluster 5, composed of a minority of FBs (53 nuclei among all the 4 samples), showed high expression of mononuclear cell differentiation genes (*Card11*, *Dock2*, *Ikzf3*) suggesting their possible belonging to an undifferentiated immune cell population.

The classification of macrophages, monocytes, precursors of T and B cells, dendritic cells, stromal cells, and T cells subclusters was performed through MyGeneSet data browser of the Immgen project (Heng et al., 2008). Both macrophage subclusters were composed of M2 macrophages (*Cd163+*, *Mrc1+*), known to be involved in anti-inflammatory and tissue repair functions (Yunna et al., 2020). M2 subcluster 1 showed characteristics of M2a (*Cd45/Ptprc, Tnf, Fn1, Cxcl16*) and M2c (*Il10* and *Tgfβ1*) subpopulations, suggesting its involvement in wound healing and remodeling. M2 subcluster 2 expressed markers of the M2c population (*Cd206/Mrc1, Cd163*, and *Mertk*), displaying immunosuppression, phagocytosis, and tissue remodeling functions (Gharavi et al., 2022; Wang et al., 2019).
